# Non-Invasive Embryo Quality Assessment via Matrix-Optimized Untargeted LC-MS Metabolomics of Spent Embryo Culture Media and Weighted Ensemble Machine Learning

**DOI:** 10.64898/2026.07.30.741666

**Authors:** Huimin Gan, Xingyao Wang, Feitai Tang, Hebatallah Ibrahim, Xuanyu Chen, Pingyuan Xie, Shuoping Zhang, Ge Lin, Jun Zeng, Hongwei Chu, Shen Zhang

## Abstract

**Background:** Non-invasive embryo quality assessment is a critical unmet need in assisted reproductive technology (ART). Preimplantation genetic testing for aneuploidy (PGT-A) is effective but requires invasive biopsy that may compromise embryo viability. Metabolomics of spent embryo culture media (SECM) offers a non-invasive alternative, yet analytical challenges—limited sample volume, high salt content, and abundant proteins—have hindered standardization and clinical translation.

**Results:** We systematically optimized sample preparation for untargeted LC-MS metabolomics of SECM using human serum as a reference. Optimal conditions were highly matrix-dependent: SECM required 7× volume of 50% acetonitrile for extraction and 40% acetonitrile for reconstitution, whereas serum required 10× volume of 100% methanol and 100% water—reflecting that SECM contains more non-polar species than serum. Applying the optimized workflow to 120 clinical SECM samples (72 euploid, 48 aneuploid), we identified 102 differential metabolites between euploid and aneuploid embryos, with prominent enrichment of lipid pathways (fatty acid metabolism, β-oxidation, sphingolipid metabolism) and involvement of amino acid (methionine, tryptophan) and TCA cycle metabolism. A weighted ensemble machine learning model discriminated aneuploid from euploid embryos with an AUC of 0.977, 100.0% specificity, and 89.6% sensitivity. Among 72 euploid embryos stratified by morphological grading (good, fair, poor), metabolic alterations progressed from mitochondrial energy deficiency (good vs. fair) to broader lipid dysregulation (fair vs. poor), with the ensemble model achieving AUCs of 0.944, 0.889, and 0.943, respectively.

**Conclusions:** This study establishes a rigorously optimized and validated SECM metabolomics workflow that overcomes key analytical barriers in this challenging matrix. Our findings demonstrate that metabolic signatures—particularly in lipid and energy metabolism—are strongly associated with both embryo ploidy and morphological quality, providing biological insights into the metabolic underpinnings of embryo developmental competence. The high predictive performance of the ensemble model supports the feasibility of non-invasive embryo assessment as a complementary tool to existing methods, with potential to reduce reliance on invasive biopsy in ART. External validation in prospective multi-center cohorts is warranted to further assess clinical utility and generalizability.

**Graphical Abstract:** 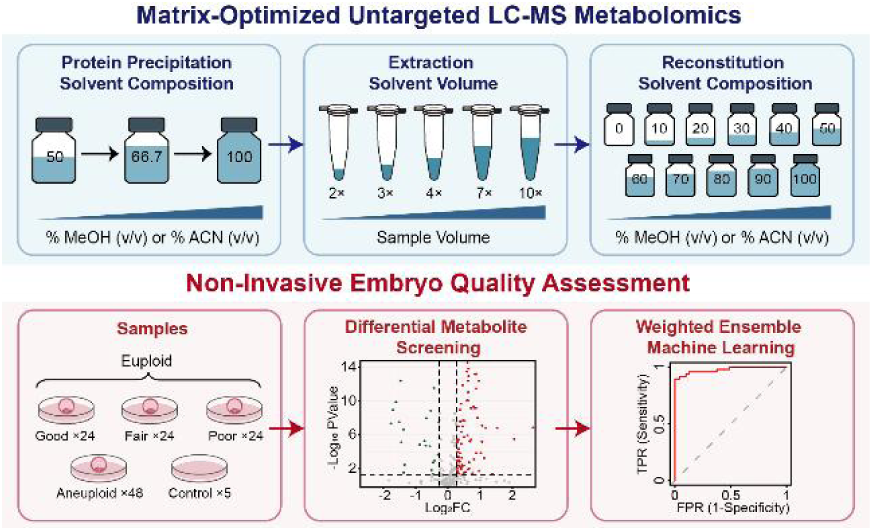

## Introduction

Since the first successful in vitro fertilization (IVF) in 1978, assisted reproductive technologies (ART) have enabled over 8 million births worldwide. Nevertheless, implantation rates remain suboptimal, ranging from 4% to 40%^1, 2^. The global trend toward elective single embryo transfer (eSET), adopted to reduce risks associated with multiple pregnancies, has increased the need for accurate, non-invasive methods to assess embryo viability and select the most competent embryo for transfer^3, 4^. Traditional embryo evaluation relies primarily on morphological assessment, including blastocyst expansion grade, inner cell mass quality, and trophectoderm appearance according to the Gardner classification system^5^. Although enhanced by time-lapse imaging and artificial intelligence-based algorithms, morphological assessment alone remains subjective and insufficient to fully capture the embryo’s developmental potential and implantation capacity^6–8^.

Preimplantation genetic testing for aneuploidy (PGT-A) is the current clinical gold standard for improving embryo selection accuracy^9, 10^. However, this invasive approach requires embryo biopsy, raising concerns about potential impacts on embryo viability and long-term developmental outcomes^11, 12^. Consequently, non-invasive alternatives that can assess embryo quality without biopsy are of growing interest, as they may provide complementary information about embryo functional status and developmental competence.

Metabolomics, the comprehensive analysis of small molecules (<1.5 kDa) in biological systems, has emerged as a powerful tool for assessing embryo quality through the analysis of SECM^13, 14^. During in vitro development, embryos continuously exchange metabolites with their surrounding environment^15^. The resulting metabolic footprint in SECM reflects the integrated activity of multiple biochemical pathways and has been correlated with embryo viability, ploidy status, and implantation potential^16^. Previous studies have identified specific metabolic signatures associated with successful pregnancy, including altered consumption of amino acids, pyruvate, and glucose^17, 18^, as well as differential release of metabolites such as alanine, glutamate, and formate^19^.

Liquid chromatography-mass spectrometry (LC-MS) has become the platform of choice for untargeted metabolomics due to its high sensitivity and wide dynamic range^20, 21^. However, applying LC-MS to SECM analysis presents unique analytical challenges. Individual embryo culture volumes are typically around 20 μL, yielding limited sample volume. Moreover, SECM contains high concentrations of inorganic salts (maintaining physiological osmolality), abundant proteins (primarily human serum albumin as a protein source), and a complex mixture of amino acids and other medium components that can cause severe matrix effects, ion suppression, and interference with metabolite detection^22, 23^. These challenges necessitate careful optimization of sample preparation protocols to maximize metabolome coverage while maintaining data quality and reproducibility^24, 25^.

Sample preparation for LC-MS metabolomics typically involves protein precipitation, metabolite extraction, and sample reconstitution prior to injection^26^. Protein precipitation is most commonly achieved using organic solvents such as methanol (MeOH) or acetonitrile (ACN), which denature and precipitate proteins while solubilizing a wide range of metabolites^27, 28^. The choice of extraction solvent critically influences extraction efficiency: MeOH generally provides broader coverage of polar metabolites, whereas ACN often performs better for lipids and non-polar compounds^29, 30^. The solvent-to-sample volume ratio must be balanced to achieve complete protein removal without excessive dilution that would reduce sensitivity for low-abundance metabolites^31^. After extraction and solvent evaporation, samples are reconstituted in a solvent compatible with subsequent chromatographic separation. The reconstitution solvent composition can substantially affect metabolite solubility, peak shape, and overall detection sensitivity^32^. Lindahl et al. demonstrated that reconstitution in 100% water maximized metabolite feature detection for methanol-extracted biological samples analyzed by reversed-phase LC-MS, challenging the common practice of using 50% organic modifier^33^. Despite the critical importance of these parameters, systematic optimization of sample preparation workflows for SECM metabolomics has been limited. Previous studies have used diverse protocols—different extraction solvents, volumes, and reconstitution conditions—complicating cross-study comparisons and hindering method standardization^13, 34^. Furthermore, the unique matrix characteristics of SECM (low protein content relative to serum but high salt and defined medium components) warrant specific optimization rather than direct adoption of protocols developed for other biofluids.

In this study, we systematically optimized key sample preparation parameters for non-targeted LC-MS metabolomics analysis of SECM, including protein precipitation solvent composition (methanol-water vs. acetonitrile-water), extraction solvent volume, and reconstitution solvent composition, using human serum as a parallel reference. Optimal conditions were highly matrix-dependent: for SECM, extraction with 7× volume of 50% ACN followed by reconstitution in 40% ACN maximized metabolome coverage, whereas serum required 10× volume of 100% MeOH for extraction and 100% water for reconstitution. Applying this SECM-optimized workflow to 120 clinical samples (72 euploid, 48 aneuploid), we identified distinct metabolic signatures associated with embryonic ploidy status. A weighted ensemble machine learning model discriminated aneuploid from euploid embryos with an AUC of 0.977. Furthermore, analysis of the 72 euploid SECM samples divided by embryo morphological grading (Good, Fair, and Poor) revealed progressive metabolic alterations in energy and lipid metabolism, with the ensemble model achieving AUCs of 0.944, 0.889, and 0.943, respectively. These findings establish a robust, non-invasive SECM metabolomics approach for embryo ploidy screening and quality assessment in ART.

## Methods

### Sample Collection and Preparation

SECM samples were collected from patients undergoing IVF treatment at the Reproductive and Genetic Hospital of CITIC-XIANGYA. Embryos were transferred to blastocyst medium, and a change in fresh medium (each embryo per 20 μl droplet) combined with complete cumulus cell removal was performed on Day 4. Blastocyst morphology was assessed on Day 5 according to the Gardner grading criteria and again on Day 6 if they failed to meet the biopsy criteria on Day 5. TE biopsy was performed if the embryo reached a morphologic grade of at least 4BC, and the corresponding SECM were collected simultaneously. For method optimization experiments, multiple SECM aliquots were pooled, mixed thoroughly, and re-aliquoted to ensure consistent starting material across all experimental conditions. Human serum samples were obtained from healthy volunteers. Blood was collected into serum separator tubes, allowed to clot for 30 min at room temperature, and centrifuged at 1500×g for 15 min at 4°C. Serum was aliquoted (50 μL per tube) and stored at −80°C until analysis. Similarly, for optimization experiments, multiple serum aliquots were pooled, mixed thoroughly, and re-aliquoted to ensure consistency. The study protocol was approved by the hospital’s Ethics Committee (Approval No. LL-SC-2023-033).

### Metabolite Extraction Procedure

Frozen SECM (20 μL) and serum (50 μL) samples were thawed on ice prior to extraction. To optimize extraction solvent composition and volume, samples were mixed with extraction solvents at ratios of 2×, 3×, 4×, 7×, and 10× the sample volume. Six extraction solvent compositions were evaluated: three methanol-water systems (100% MeOH, 66.7% MeOH, 50% MeOH, v/v) and three acetonitrile-water systems (100% ACN, 66.7% ACN, 50% ACN, v/v). For ternary solvent evaluation, a mixture of 40% MeOH-40% ACN-20% H₂O (v/v/v) was tested at 3× and 7× sample volumes for SECM, and 3× and 10× sample volumes for serum. Extraction was performed by adding ice-cold extraction solvent to each sample, followed by vigorous vortexing for 30 s. Samples were incubated at −20°C for 20 min to enhance protein precipitation and then centrifuged at 21,300×g for 15 min at 4°C. The supernatant was carefully transferred to a new pre-chilled microcentrifuge tube and evaporated to dryness using a SpeedVac vacuum concentrator (Thermo Fisher Scientific, Waltham, MA, USA) at 4°C. Dried extracts were stored at −80°C until reconstitution and LC-MS analysis. All extraction conditions were tested in triplicate.

### Optimization of Reconstitution Conditions

Following extraction optimization, samples were processed using the optimal extraction conditions for each matrix: for SECM, extraction with 7× volume of 50% ACN; for serum, extraction with 10× volume of 100% MeOH. Dried extracts were reconstituted in 25 μL of eleven different reconstitution solvent compositions, prepared by mixing ACN (for SECM) or MeOH (for serum) with water in 10% increments: 100%, 90%, 80%, 70%, 60%, 50%, 40%, 30%, 20%, 10%, and 0% organic content. Samples were vortexed for 30 s and centrifuged at 15,000×g for 15 min at 4°C to remove any insoluble material. Each reconstitution condition was evaluated in triplicate.

### LC-MS/MS Analysis

Non-targeted metabolomics analysis was performed using a Vanquish Flex UHPLC system coupled to an Orbitrap Exploris 120 mass spectrometer (Thermo Fisher Scientific, Bremen, Germany). Details for chromatographic separation and MS analysis were described in Supporting Information.

### Data Processing and Statistical Analysis

Raw LC-MS data were processed using Compound Discoverer 3.3 software (Thermo Fisher Scientific). Details for data processing and statistical analysis were described in Supporting Information.

### Machine learning model construction

All machine learning procedures were carried out in RStudio (version 4.5.1). A leave-one-out cross-validation (LOOCV) strategy was adopted for model training and evaluation. Two separate prediction models were developed: one for aneuploidy prediction and one for embryo morphological quality grading. Candidate biomarkers were preselected based on univariate and multivariate statistical criteria as well as analytical stability. For the aneuploidy prediction model, features with |FC| > 1.2, p < 0.05, and a CV in QC samples < 0.25 were retained. For the morphological quality grading model, the threshold was set at CV < 0.15 (with |FC| > 1.2 and p < 0.05). Metabolite data were then normalized by Z-score to eliminate scale differences.

Seven base machine learning models were constructed: logistic regression, linear discriminant analysis (LDA), support vector machine (SVM), random forest, k-nearest neighbors (KNN), decision tree, and naive Bayes. Each model was trained under the LOOCV framework: in each iteration, one sample was held out as the test set and the remaining samples were used for training. The process was repeated until every sample had been predicted once. Performance metrics for each model included accuracy, sensitivity, specificity, positive predictive value (PPV), negative predictive value (NPV), area under the receiver operating characteristic curve (AUC), and F1 score. Feature importance was calculated using the random forest model based on the mean decrease in Gini impurity, and candidate biomarkers were ranked accordingly.

Finally, a weighted voting ensemble model was constructed by assigning weights based on the prediction accuracy of each base model; only models with accuracy > 0.5 were included. For the aneuploidy prediction model, the optimal classification threshold was determined by maximizing the Youden index^35^, and samples were classified as either “euploid” or “aneuploid”. For the morphological quality grading model, the one-vs-rest (OvR) strategy was employed to decompose the three-class problem (good-, fair-, and poor-quality embryos) into multiple binary classification tasks. Performance metrics for each class (sensitivity, specificity, PPV, NPV, F1 score, and AUC) were calculated, and the final predicted class was determined as the one with the highest weighted probability.

## Results

### Optimization of Metabolite Extraction Solvent Composition and Volume for SECM and Serum

The choice of metabolite extraction solvent fundamentally determines the range of metabolites extracted from biological samples, as solvent polarity directly affects the solubility of different metabolite classes^29, 30^. To investigate the effects of extraction solvent on metabolome coverage, quantitative reproducibility, and relative recovery, we systematically evaluated six extraction solvents comprising methanol-water and acetonitrile-water systems at three organic compositions (100%, 66.7%, and 50%), each tested at five solvent-to-sample volume ratios (2×, 3×, 4×, 7×, and 10×) (**Figure 1A**). As shown in **Figure 1B** and **Table S1**, acetonitrile-based solvents generally outperformed methanol-based solvents in total metabolite identifications for SECM. Among methanol-based solvents, the number of identified metabolites increased as the MeOH proportion decreased, reaching a maximum with 10× volume of 50% MeOH, which yielded an average of 188 metabolites across three replicates, and identifications decreased substantially at lower water ratio (66.7% MeOH: 167 ± 4; 100% MeOH: 138 ± 2). Among acetonitrile-based solvents, 50% ACN at 7× volume achieved a maximum of 194 metabolites on average; and the identifications also decreased at lower water ratio. Overall, in most cases, increasing the water ratio in the extraction solvent enhanced metabolite identification for both systems.

**Figure 1.**
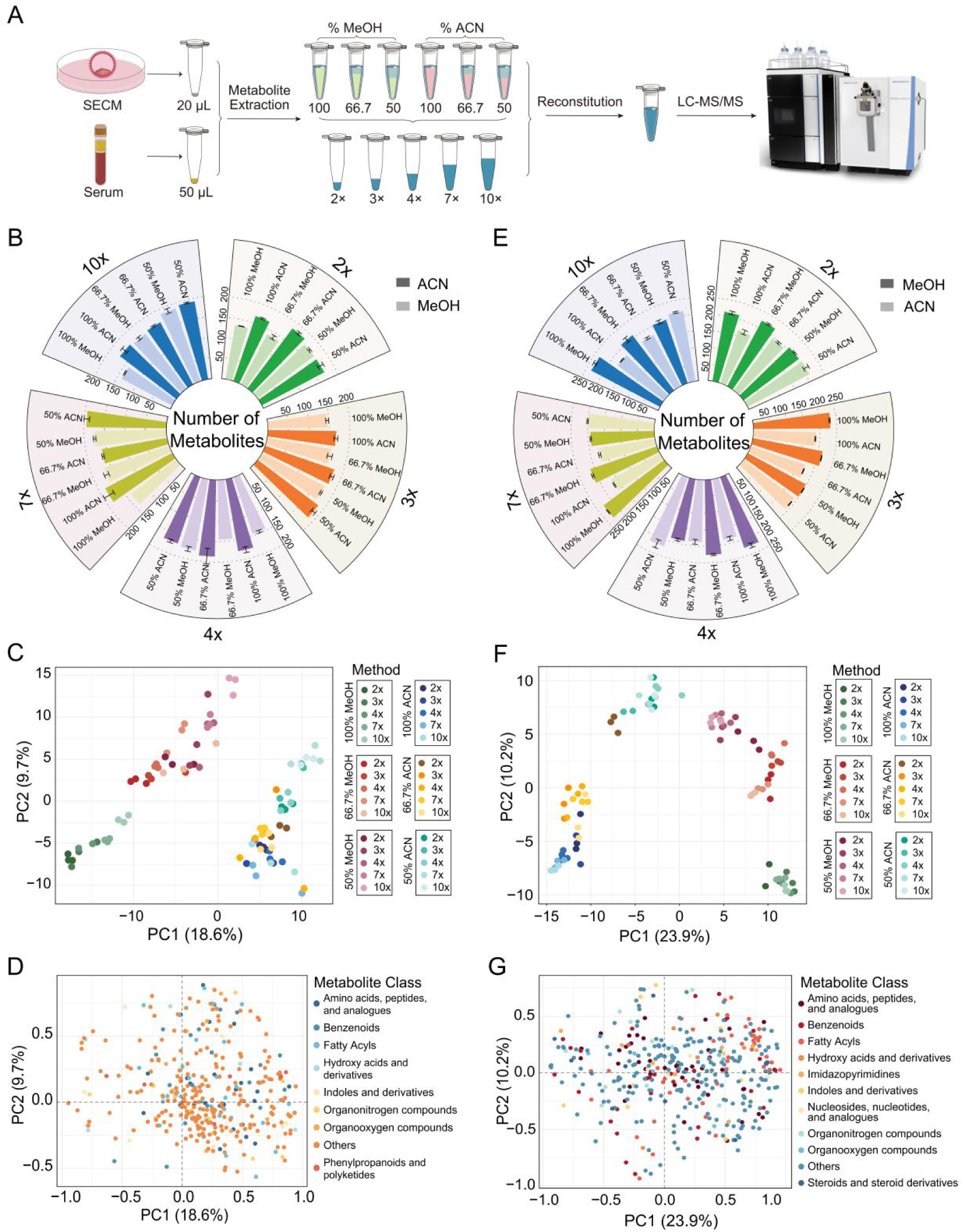
Optimization of the metabolite extraction methods for SECM and serum. (A) The optimization workflow of metabolite extraction for SECM and serum. (B) Number of identified metabolites from SECM by different extraction solvent composition and volume. (C) PCA score scatter plot of peak area response for the different extraction solvent composition and volume for SECM. (D) The corresponding loading plot of different metabolite class for Figure 1C. (E) Number of identified metabolites from serum by different extraction solvent composition and volume. (F) PCA score scatter plot of peak area response for the different extraction solvent composition and volume for serum. (G) The corresponding loading plot of different metabolite class for Figure 1F.

Extraction solvent volume had a relatively minor effect, though larger volumes modestly improved the number of identified metabolites. PCA clearly distinguished the methanol-based solvents from acetonitrile-based solvents, with samples separated along the central axis (**Figure 1C**). Methanol-based solvents extract clustered on the left, while acetonitrile–based solvents grouped on the right, showing trends associated with extraction solvent composition and volume, indicating good intergroup differentiation. The corresponding loading plot revealed that most metabolite classes were located on the right side, further supporting that the acetonitrile–based solvents extracted a broader range of metabolite classes more effectively in SECM (**Figure 1D**). The coefficient of variation (CV) was calculated for all metabolites identified by each method. The median CV for all methods was below 0.2, indicating satisfactory quantitative reproducibility (**Figure S1**). Among these, the method with the highest metabolite number in the acetonitrile–based solvents (7× volume of 50% ACN) had a median CV of 0.17. Relative recovery of metabolites was also evaluated using this solvent as a reference. For each other method, metabolite intensities were normalized to the mean intensity of the corresponding metabolite in the reference, and normalized values were averaged within each metabolite class. Relative recovery analysis showed that while some metabolite classes exhibited no clear dependence on the extraction method, four classes—amino acids, peptides, and analogues; benzenoids; fatty acyls; and organooxygen compounds—displayed notably higher intensities with the 7× volume of 50% ACN method (**Figure S2**), further confirming its superior extraction performance.

For serum samples, a trend opposite to that observed in SECM was found: the acetonitrile-based solvents generally identified fewer metabolites than the methanol-based solvents, indicating markedly inferior extraction efficiency for serum metabolites (**Figure 1E**, **Table S2**). Among methanol-based solvents, the number of identified metabolites decreased as the water ratio increased, reaching a maximum of 234 metabolites with 10× volume of 100% MeOH, consistent with published results^33^. Among acetonitrile-based solvents, metabolite identification increased with water ratio, peaking at 4× volume of 50% ACN with an average of 220 metabolites. Extraction solvent volume had a relatively minor effect, though larger volumes modestly improved metabolite identification. Similar to SECM, PCA clearly distinguished methanol-based and acetonitrile-based solvents, with samples symmetrically separated along the axis (**Figure 1F**). The corresponding loading plot revealed that most metabolite classes were distributed on both sides; notably, a greater number of fatty acyls were located on the right side, suggesting that this compound class was more effectively extracted by methanol-based solvents (**Figure 1G**). The median CV for all methods was below 0.2, indicating satisfactory quantitative reproducibility, with the methanol-water system mostly exhibiting lower CV values than the acetonitrile-water system (**Figure S3**). Relative recovery was evaluated using the 10× volume of 100% MeOH extractant as a reference. Relative recovery analysis showed that two classes—fatty acyls and hydroxy acids and derivatives—displayed notably higher intensities with the 10× volume of 100% MeOH method, further confirming its superior extraction performance for these compound classes (**Figure S4**).

The marked difference in optimal extraction conditions between SECM and serum prompted us to investigate whether a ternary solvent system combining MeOH and ACN might offer complementary benefits for metabolite extraction. A well optimized and widely used ternary solvent system in metabolomics studies, 40% ACN-40% MeOH-20% H₂O (AMW20), was adopted to compare its performance against the respective optimal binary solvents identified above (7× 50% ACN for SECM; 10× 100% MeOH for serum). For SECM, the ternary solvent systems (3× and 7×) identified 182 ± 5 and 173 ± 4 metabolites, respectively - significantly fewer than the 203 ± 5 achieved with 7× 50% ACN (p < 0.005; **Figure S5A**). Venn diagram analysis revealed that 82% of metabolites identified by ternary extraction were also detected by the optimal binary method, with only 5-8% unique to ternary extraction (**Figure S5B**). The CV values were comparable across all methods (median 0.10-0.11), indicating the good reproducibility of all these methods (**Figure S5C**). For serum, ternary solvents (3× and 10×) identified 187 ± 1 and 245 ± 6 metabolites, both fewer than the 250 ± 4 achieved with 10× 100% MeOH (**Figure S5D**). Overlap still revealed that most of metabolites identified by ternary extraction were also detected by the optimal binary method (**Figure S5E**). The CV values were also comparable across all methods (median 0.09-0.10) (**Figure S5F**). These results demonstrate that ternary solvent mixtures, despite containing both MeOH and ACN, do not provide the broad metabolite coverage that might be expected from combining the two organic solvents. This finding likely reflects the fact that solvent mixtures create a homogeneous polarity environment rather than the selective extraction that occurs when using pure solvents alone^36^.

### Comparative Metabolomics of SECM and Serum

The above results suggest that optimal sample preparation conditions are fundamentally matrix-dependent: for SECM, 50% ACN at 7× sample volume provided maximum metabolome coverage, whereas for serum, pure MeOH at 10× sample volume yielded the best identification depth. To further investigate compositional differences between SECM and serum, we compared all metabolites identified by all methods in this study for the two sample types (**Figure 2A**). Among 391 metabolites identified in SECM and 396 in serum, only 174 (∼44%) were common to both matrices, indicating substantial differences in chemical composition. Comparison of logP distributions revealed that more polar compounds (logP < 0) were detected in serum, while more non-polar species (logP > 0) were identified in SECM (**Figure 2B**). This observation is consistent with the distinct biological origins of the two matrices. Serum is a complex biofluid rich in circulating polar metabolites—such as amino acids, organic acids, and carbohydrates—that participate in systemic metabolism. In contrast, SECM reflects the metabolic activity and secretory products of the embryo, including lipid-derived signaling molecules and membrane-associated metabolites, which likely contribute to a higher proportion of non-polar or amphipathic compounds. When the identified metabolites were classified according to the HMDB database, both matrices were dominated by fatty acyls, benzenoids, amino acids and peptides, organoheterocyclic compounds, and organic acids and derivatives (**Figure 2C**). SECM samples contained a larger proportion of unclassified compounds compared to serum, followed by fatty acyls (11.76%), benzenoids (7.67%), and amino acids (6.91%). Serum samples exhibited a higher relative abundance of polar metabolite classes, including fatty acyls (13.89%), amino acids (10.61%), and benzenoids (8.33%). These differences reflect the distinct physicochemical properties of the two matrices and underscore a critical principle: sample preparation protocols should not be transferred directly between different sample types without systematic re-optimization. This principle has important implications for the growing field of SECM metabolomics. Many published studies have adopted protocols originally developed for serum or plasma analysis, potentially compromising sensitivity and coverage for this unique matrix. Our optimized SECM protocol offers a useful reference for future biomarker discovery studies, as it improves the detection of potentially informative metabolites while maintaining good analytical reproducibility.

**Figure 2.**
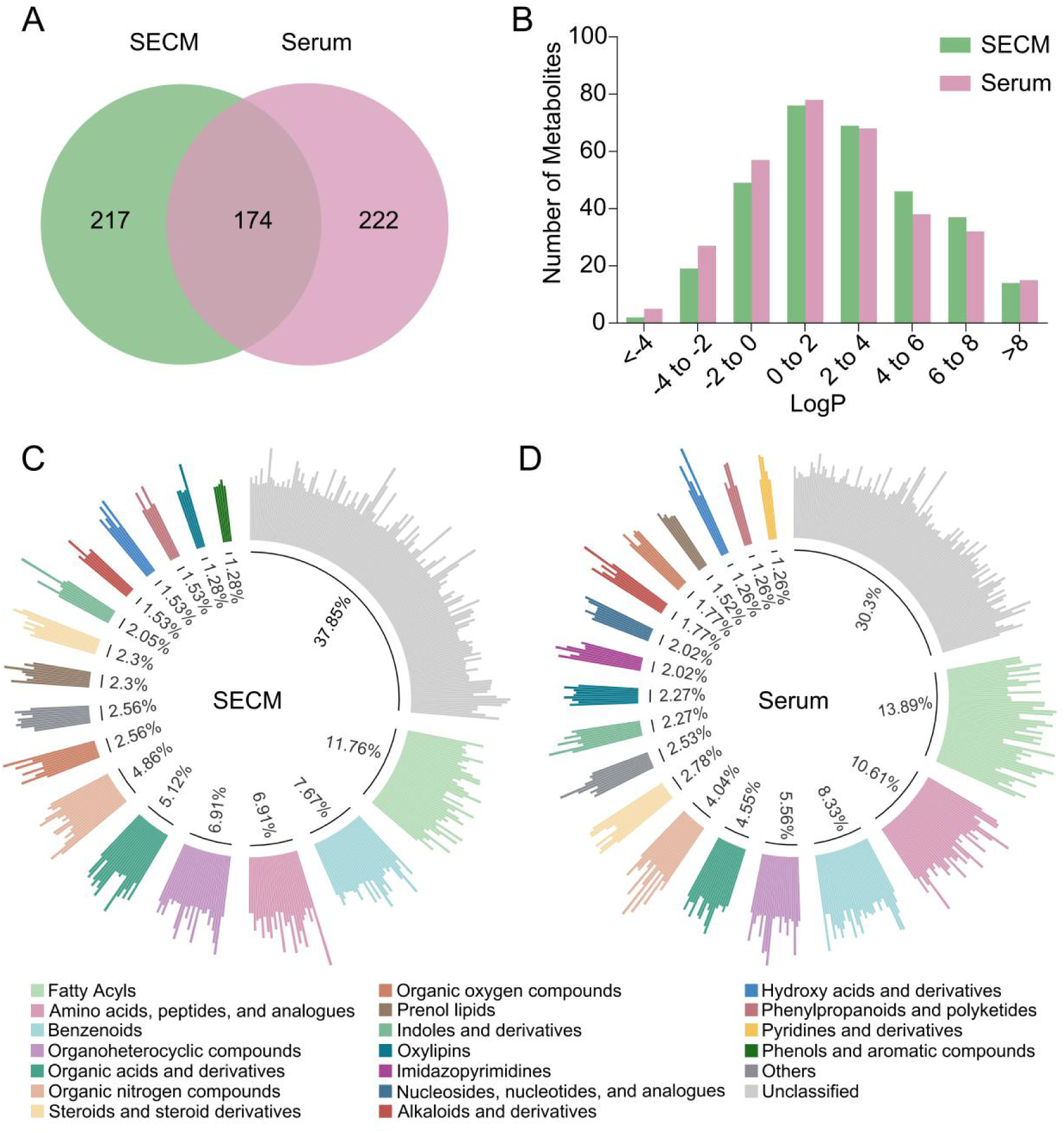
Comparison of the metabolites in SECM and serum. (A) The overlap of all the metabolites identified from SECM and serum. (B) The log P distribution of all metabolites identified in SECM and serum as listed in the Human Metabolome Database (HMDB). (C) Metabolites classification proportion statistics in SECM. (D) Metabolites classification proportion statistics in serum. The unclassified categories grouped as “Unclassified” and classes containing fewer than five entries grouped as “Others”.

### Optimization of Reconstitution Solvent Composition

The reconstitution step, although often overlooked in method development, critically influences metabolite solubility, chromatographic behavior, and overall detection sensitivity. Using the optimal extraction conditions for each matrix (7× 50% ACN for SECM; 10× 100% MeOH for serum), we systematically evaluated eleven reconstitution solvent compositions ranging from 0% to 100% organic content (**Figure 3A**). For SECM samples, reconstitution solvent composition dramatically affected metabolome coverage (**Figure 3B**). Maximum metabolite identifications (195 ± 14) were achieved with 40% ACN in water. The poor performance of 100% ACN likely results from two factors: first, highly polar metabolites may precipitate or adsorb to vial surfaces when reconstituted in predominantly organic solvent; second, injection of samples in strong solvent can cause poor peak focusing, particularly for early-eluting polar compounds. PCA of reconstitution conditions (**Figure 3C**) revealed a clear trajectory along PC1 corresponding to increasing water content, demonstrating systematic and reproducible effects of reconstitution solvent composition on the detected metabolome. Reproducibility was satisfactory across all conditions (median CV range: 0.08-0.25) (**Figure 3D**). Relative recovery analysis using 40% ACN as reference showed that this composition provided superior recovery for benzenoids, fatty acyls, organic phosphoric acids, steroids/steroid derivatives, compared to other compositions (**Figure S6**). For serum samples, reconstitution solvent optimization revealed a strikingly different pattern (**Figure 3E**). Metabolite identifications decreased progressively as MeOH content increased from 0% to 80%, then increased slightly at higher organic content. Maximum identifications (239 ± 2) were achieved with 100% water, with 80% MeOH yielding significantly fewer identifications (212 ± 3). The slight increase in identifications at very high organic content (>80%) may correspond to recovery of a small number of highly hydrophobic metabolites that are poorly soluble in water, but the overall trend strongly favors aqueous reconstitution for maximizing coverage of the serum metabolome. These findings align with those of Lindahl et al., who reported that 100% water maximized feature detection for methanol-extracted serum samples analyzed by reversed-phase LC-MS. PCA showed clear separation according to water content (**Figure 3F**). Reproducibility was excellent across all conditions (median CV: 0.05-0.08), with pure water achieving a median CV of 0.08 (**Figure 3G**). Relative recovery analysis confirmed that pure water provided superior recovery for amino acids/peptides, hydroxy acids, indoles/derivatives, and imidazopyrimidines (**Figure S7**).

**Figure 3.**
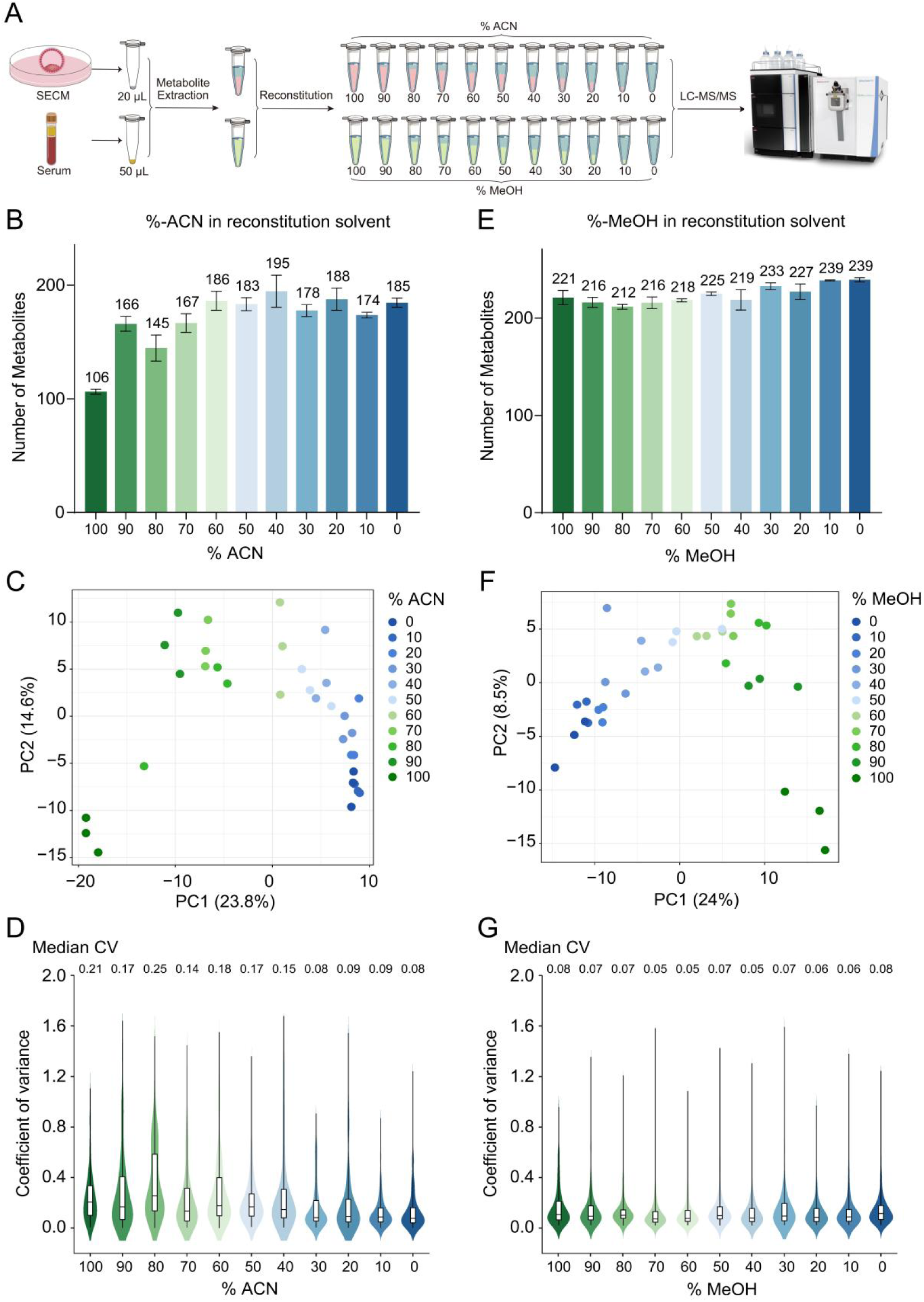
Optimization of reconstitution solvent composition for SECM and serum. (A) The optimization workflow of metabolite reconstitution solvent composition for SECM and serum. (B) Number of identified metabolites from SECM by different ratio of ACN. (C) PCA score scatter plot of peak area response for the different reconstitution solvent composition for SECM. (D) The coefficient of variation (CV) distribution of peak area across replicates by different reconstitution solvent composition for SECM. (E) Number of identified metabolites from serum by different ratio of MeOH. (F) PCA score scatter plot of peak area response for the different reconstitution solvent composition for serum. (G) The coefficient of variation (CV) distribution of peak area across replicates by different reconstitution solvent composition for serum.

In addition to sample preparation optimization, we compared profile and centroid MS acquisition modes using SECM samples processed with the optimized protocol (7× 50% ACN extraction, reconstitution in 40% ACN or 50% ACN). Centroid mode consistently identified more metabolites than profile mode under both reconstitution conditions (**Figure S8A**). For 40% ACN reconstitution, centroid mode identifed 209 ± 6 metabolites, whereas profile mode only yielded 196 ± 4 metabolites (*p-*value = 0.0013); for 50% ACN, centroid mode identified 187 ± 5 metabolites, significantly more than the 171 ± 2 metabolites identified by profile mode (*p-*value = 0.0024). Notably, the majority of metabolites were detectable under both acquisition modes (**Figure S8B**). Reproducibility was also superior in centroid mode, with median CV values of 0.06 and 0.03 compared to 0.11 and 0.16 for profile mode (**Figure S8C**). The improved performance of centroid mode is likely due to its real-time peak detection and recording of only the peak apex intensity with accurate mass, which effectively filters high-frequency noise and baseline fluctuations, yielding cleaner spectra with higher signal-to-noise ratios. This approach enhances feature detection, alignment, and integration for improved quantification—particularly for low-abundance metabolites—and is inherently compatible with spectral library search algorithms that are predominantly based on centroid spectra, thus enabling more reliable identifications.

### Untargeted Metabolomics of SECM Identifies Biomarkers for Aneuploidy Prediction

Utilizing the optimized sample preparation workflow, we performed untargeted metabolomics analysis on 120 SECM samples (72 euploid, 48 aneuploid). The biopsy results of the blastocysts corresponding to these SECM were shown in **Table S3**. Five blank culture medium samples were included as background controls. A total of 312 metabolites were quantified. PLS-DA revealed a clear separation between euploid and aneuploid groups (**Figure 4A**). Comparison between euploid and aneuploid groups identified 102 differential expressed metabolites (DEMs, p < 0.05, |FC|>1.2), of which 80 were upregulated and 22 downregulated in the aneuploid group (**Figure 4B**). Classification according to the HMDB database showed that the identified metabolites mainly classified into amino acids, fatty acids, organic acids. Notably, a large number of DEMs were fatty acids (**Figure 4C**), including polyunsaturated fatty acids, saturated fatty acids, fatty acid amides, and lipid peroxidation products. Among these, docosahexaenoic acid, oleic acid, and docosapentaenoic acid have been closely linked to membrane stability, mitochondrial function, and embryonic developmental potential^37, 38^. Conversely, elevated levels of saturated fatty acids such as palmitic acid and stearic acid may induce lipotoxicity, oxidative stress, and apoptosis, thereby impairing blastocyst formation^39^. Among amino acids, methionine is a crucial substrate for one-carbon metabolism; it participates in DNA and histone methylation via S-adenosylmethionine (SAM), playing an essential role in epigenetic regulation and cell differentiation. Abnormal methionine metabolism may compromise blastocyst formation and developmental potential (**Figure 4D**)^40, 41^. Regarding organic acids, the identified DEMs are involved in energy metabolism, the tricarboxylic acid (TCA) cycle, and amino acid metabolism (**Figure 4E**). Pyruvic acid and L-(+)-lactic acid are among the most important energy substrates for early embryo development; during the cleavage stage, embryos primarily rely on pyruvate and lactate for energy, and their uptake and metabolic levels correlate with developmental potential and blastocyst formation^42, 43^. Citric acid and isocitric acid, as key TCA cycle intermediates, reflect mitochondrial oxidative phosphorylation and energy status, and mitochondrial function is considered a critical determinant of embryo quality^44, 45^. Pathway enrichment analysis of these DEMs revealed significant enrichment of multiple lipid-related pathways, including α-linolenic acid and linoleic acid metabolism, arachidonic acid metabolism, fatty acid biosynthesis and elongation, mitochondrial β-oxidation, sphingolipid metabolism, and steroidogenesis (**Figure 4F**). These pathways involve key fatty acids and their derivatives, such as adrenic acid, arachidonic acid, docosahexaenoic acid (DHA), docosapentaenoic acid (DPA), palmitic acid, and stearic acid, indicating a central regulatory role of lipid metabolism in embryo development. Early embryonic development is highly dependent on fatty acid metabolism for energy; in particular, mitochondrial β-oxidation provides a critical ATP source for cell division and differentiation. Moreover, polyunsaturated fatty acids (e.g., DHA and adrenic acid) not only contribute to membrane architecture but also act as signaling molecules regulating cell proliferation, inflammatory responses, and redox homeostasis. Sphingolipid metabolism and steroidogenesis are closely associated with apoptosis, membrane stability, and the hormonal microenvironment, while bile acid synthesis and glycerolipid metabolism may reflect material exchange and metabolic adaptation between the embryo and its culture environment. Notably, enrichment of propionate metabolism, branched-chain amino acid degradation (valine, leucine, and isoleucine degradation), and sulfate metabolism further suggests that embryos undergo not only lipid-dependent energy production but also amino acid metabolic reprogramming and redox regulation. The coordinated changes in these metabolic networks may jointly influence embryonic developmental potential and chromosomal stability.

**Figure 4.**
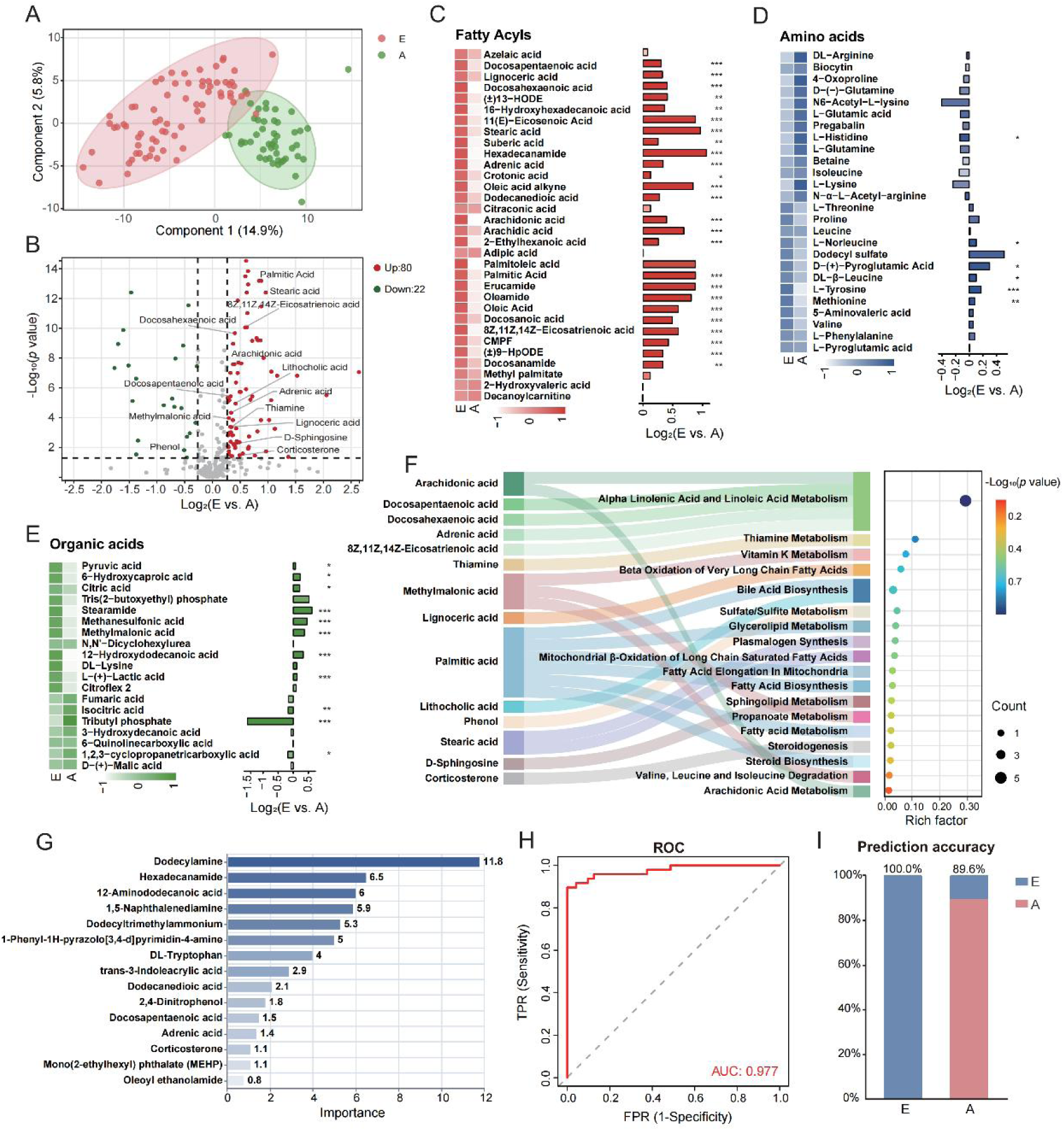
Untargeted metabolomics of SECM from euploid (E) and aneuploid (A) embryos. (A) PLS-DA analysis of metabolomic data of SECM from E and A groups. (B) Volcano plot showing differential expressed metabolites (DEMs) between E and A groups. Metabolites upregulated in group E are shown in red, downregulated in green. (C-E) Heatmaps of relative concentrations (Z-score) and fold changes for identified metabolites (fatty acids, amino acids, organic acids). *p < 0.05, **p < 0.01, ***p < 0.001 (Student’s t-test). Z-score normalization was applied to better display the relative concentrations of the mean values of each metabolite between the two groups. (F) Pathway enrichment analysis of differential metabolites between E and A groups. (G) Mean decrease in accuracy from random forest analysis to identify potential biomarkers. This metric indicates the reduction in prediction accuracy when the values of a variable are permuted; a larger reduction indicates greater importance of the variable. (H) ROC curves evaluating the performance of the biomarker model in discriminating the two groups. (I) Prediction accuracy of the classification model in distinguishing the two groups.

To establish a predictive model for aneuploidy based on the metabolomics results, we employed a leave-one-out cross-validation (LOOCV) strategy for model training and evaluation on these 120 samples. Fifteen candidate biomarkers were preselected based on univariate and multivariate statistical criteria and analytical stability (**Figure S9, Table S4**). These candidate biomarkers included lipid-related metabolites such as docosapentaenoic acid and adrenic acid, suggesting pronounced lipid metabolic reprogramming and membrane lipid remodeling^38, 46^. Abnormal levels of dodecanedioic acid and 2,4-dinitrophenol pointed to altered mitochondrial β-oxidation and energy metabolism^47, 48^. In addition, changes in DL-tryptophan and trans-3-indoleacrylic acid indicated the involvement of oxidative stress and immune homeostasis regulation in embryo development^18, 49^. Performance metrics for each model, including accuracy, sensitivity, specificity, PPV, NPV, AUC, and F1 score, are summarized in **Table S5**. Feature importance was calculated using the random forest model based on the mean decrease in Gini impurity, and candidate biomarkers were ranked accordingly (**Figure 4G**). Receiver operating characteristic (ROC) curve analysis yielded an area under the curve (AUC) of 0.977 for discriminating aneuploid from euploid embryos (**Figure 4H**). Using the Youden index to determine the optimal probability threshold (0.6422), the model achieved prediction accuracies of 100.0% for euploid and 89.6% for aneuploid embryos (**Figure 4I**). These results demonstrate that the predictive model can accurately distinguish between euploid and aneuploid samples, indicating excellent discriminatory performance and potential for clinical application.

### Untargeted Metabolomics of SECM from Embryos with Different Morphological Grade Identifies Biomarkers for Embryo Quality Assessment

The 72 euploid SECM samples were further divided into good-quality (G), fair-quality (F), and poor-quality (P) groups according to the morphological grades of the embryos, with 24 samples per group (**Table S3**). We performed untargeted metabolomics analysis on these three groups to identify metabolites associated with embryo quality. PLS-DA revealed significant differences among the G, F, and P groups (**Figure 5A**). Pairwise comparisons identified 65 DEMs (p < 0.05, |FC|>1.2) between G and F, of which 48 were also differentially expressed between G and P (**Figure 5B, C**). Between G and P, 106 DEMs were identified, with 56 also differentially expressed between F and P. Between F and P, 92 DEMs were identified, with 33 also differentially expressed between G and F. The substantial overlap in DEMs among the three pairwise comparisons suggests common metabolic alteration features across different quality grades, and these shared metabolites may participate in similar biological regulation mechanisms during embryo growth and development. Notably, amino acids, fatty acids, and organic acids all exhibited pronounced differences among the three groups (**Figure S10**). Pathway enrichment analysis of DEMs from different morphological grades revealed significant metabolic remodeling across all three groups, primarily involving three core metabolic networks: energy metabolism pathways (citric acid cycle, Warburg effect, transfer of acetyl groups into mitochondria), lipid metabolism pathways (fatty acid metabolism, fatty acid biosynthesis, β-oxidation of very long chain fatty acids, plasmalogen synthesis, sphingolipid metabolism), and amino acid and redox regulation pathways (tryptophan metabolism, phenylalanine and tyrosine metabolism, nicotinate and nicotinamide metabolism). Comparison between G and F groups showed significant changes in citric acid, isocitric acid, L-tryptophan, and D-sphingosine, suggesting that the main differences between high- and medium-quality embryos lie in TCA cycle activity, mitochondrial energy conversion efficiency, and tryptophan and sphingolipid metabolism (**Figure 5D**). Previous studies have shown that enhanced TCA cycle activity supports rapid embryonic cell proliferation, while sphingolipid metabolism regulates apoptosis and membrane integrity, both critical for blastocyst formation^50–52^. In the G vs. P comparison, in addition to citric acid and isocitric acid, changes were observed in nicotinamide, proline, and several long-chain fatty acids (docosapentaenoic acid, palmitic acid, stearic acid) (**Figure 5E**). This comparison enriched a broader set of pathways, including fatty acid β-oxidation, nicotinamide metabolism, and branched-chain amino acid degradation, indicating deeper metabolic differences between high- and low-quality embryos, particularly reflected in mitochondrial oxidative phosphorylation capacity, NAD⁺-dependent redox homeostasis, and fatty acid energy supply efficiency. The F vs. P comparison mainly involved lipid metabolites such as palmitic acid, lignoceric acid, 8Z,11Z,14Z-eicosatrienoic acid, and deoxycholic acid (**Figure 5F**), with enrichment in fatty acid elongation, bile acid biosynthesis, and steroid metabolism pathways. This suggests that differences between medium- and low-quality embryos are more related to lipid metabolism regulation and membrane lipid composition, whereas the degree of energy metabolism disturbance is relatively milder. Overall, all three pairwise comparisons commonly enriched the citric acid cycle, Warburg effect, fatty acid β-oxidation, and lipid biosynthesis pathways, indicating that mitochondrial energy metabolism and lipid metabolic imbalance may be common mechanisms affecting embryonic developmental potential. The additional involvement of nicotinamide, proline, and branched-chain amino acid metabolism in the high- vs. low-quality comparison further suggests that as embryo quality declines, metabolic disturbances expand from simple energy supply deficiencies to redox imbalance and amino acid metabolic reprogramming. Finally, a correlation heatmap demonstrated that the expression of these pathway-enriched differential metabolites varied significantly across different morphological grades (**Figure 5G**).

**Figure 5.**
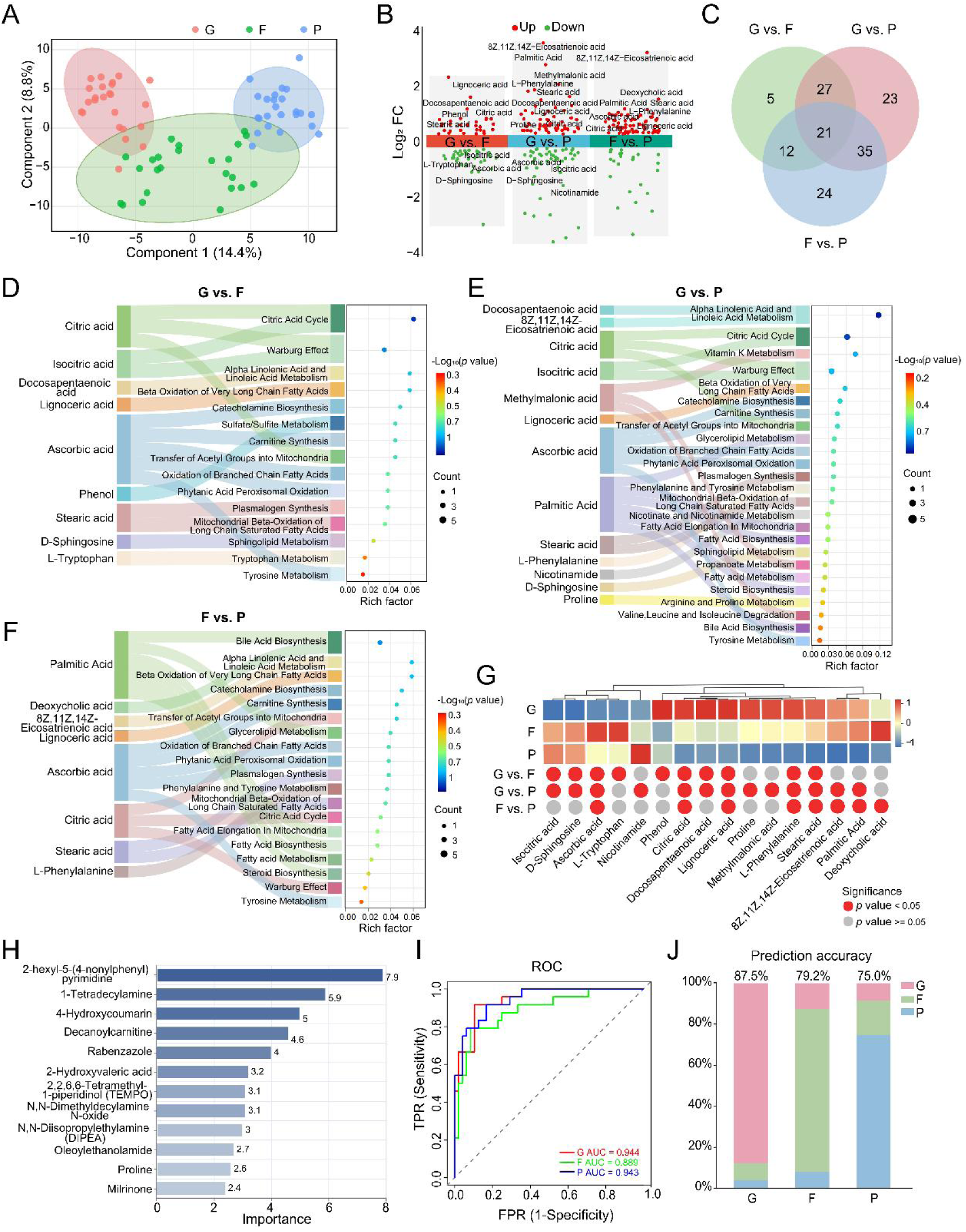
Untargeted metabolomics of SECM from embryos with different morphological grade. (A) PLS-DA analysis of metabolomic data from SECM of different morphologically graded embryos: good (G), fair (F), and poor (P) quality. (B) Volcano plots showing pairwise fold changes in metabolite abundance among the G, F, and P groups. Metabolites with upregulated abundance are shown in red, and downregulated ones in green. (C) Venn diagram illustrating the overlap in the number of differential metabolites identified from pairwise comparisons between groups. (D–F) Pathway enrichment analyses of differential metabolites from pairwise group comparisons. (G) Heatmap of differential metabolites identified from the enrichment analyses across group comparisons. (H) Mean decrease in accuracy from random forest analysis to identify potential biomarkers. This metric indicates the reduction in prediction accuracy when the values of a variable are permuted; a larger reduction signifies greater importance of the variable. (I) ROC curves evaluating the performance of the biomarker model in discriminating one population from the others. (J) Prediction accuracy of the classification model in distinguishing among the three groups.

To establish a predictive model for embryo quality grading based on SECM metabolomics, we employed a LOOCV strategy for model training and evaluation on 72 samples. Twelve candidate biomarkers were preselected based on univariate and multivariate statistical criteria and analytical stability (**Figure S11, Table S6**). Changes in decanoylcarnitine and oleoyl ethanolamide suggested altered fatty acid oxidation and mitochondrial energy metabolism during embryo development^53, 54^. Proline and 2,2,6,6-tetramethyl-1-piperidinol (TEMPO) indicated the involvement of oxidative stress and redox homeostasis regulation^55, 56^. In addition, changes in milrinone pointed to the potential influence of cAMP signaling and cell cycle regulation on early embryonic developmental potential^57, 58^. Performance metrics for each model, including overall accuracy, Kappa coefficient, macro-average sensitivity, specificity, PPV, NPV, F1 score, and AUC, are summarized in **Table S7**. Class-specific performance metrics (sensitivity, specificity, PPV, NPV, F1 score, AUC) are provided in **Table S8**. Feature importance was calculated using the random forest model based on the mean decrease in Gini impurity, and candidate biomarkers were ranked accordingly (**Figure 5H**). ROC curve analysis yielded AUCs of 0.944 (G vs. F and P), 0.889 (F vs. G and P), and 0.943 (P vs. G and F), indicating excellent discriminative ability for G and P and good discriminative ability for F (**Figure 5I**). The prediction accuracies for G, F, and P were 87.5%, 79.2%, and 75.0%, respectively, demonstrating that the predictive model can effectively distinguish among different embryo quality grades (**Figure 5J**).

## Discussion

In this study, we systematically optimized the sample preparation workflow for untargeted LC-MS metabolomics analysis of SECM and demonstrated its clinical utility for non-invasive embryo quality assessment. Our findings establish several important principles for SECM metabolomics and provide a robust methodological foundation for future biomarker discovery and clinical translation.

### Matrix-Dependent Optimization of Sample Preparation

A key finding of this study is that optimal sample preparation conditions are fundamentally matrix-dependent. For SECM, extraction with 7× volume of 50% ACN maximized metabolite identifications (194 metabolites), whereas serum required 10× volume of 100% MeOH (234 metabolites). This divergence is not merely technical but reflects profound compositional differences between the two biofluids. SECM contains a higher proportion of non-polar and amphipathic compounds (logP > 0), consistent with its origin as a secretory product of embryos actively engaged in membrane synthesis and lipid signaling. In contrast, serum is enriched in polar circulating metabolites such as amino acids, organic acids, and carbohydrates that participate in systemic metabolism. This compositional distinction explains why ACN-based solvents, which are more effective for extracting lipids and less polar compounds, performed better for SECM, whereas MeOH, which provides broader coverage of polar metabolites, was superior for serum.

The reconstitution solvent composition similarly exhibited matrix-specific effects. For SECM, 40% ACN maximized metabolite coverage, whereas for serum, 100% water was optimal. The superiority of 40% ACN for SECM likely reflects improved solubility of the more hydrophobic metabolites present in this matrix, as well as better chromatographic peak focusing for early-eluting compounds. For serum, aqueous reconstitution aligns with the polar nature of its predominant metabolites and minimizes precipitation of hydrophilic species. These findings corroborate and extend the work of Lindahl et al., who reported that 100% water maximized feature detection for methanol-extracted serum samples^33^. However, our results demonstrate that this “aqueous reconstitution” rule cannot be generalized to all biofluids, particularly those with distinct physicochemical properties such as SECM.

The ternary solvent system (40% ACN-40% MeOH-20% H₂O), despite containing both organic solvents, did not outperform the optimal binary systems for either matrix. This result may seem counterintuitive, as combining MeOH and ACN might be expected to capture both polar and non-polar metabolites. However, solvent mixtures create a homogeneous polarity environment rather than the selective extraction achieved with pure or predominantly single-solvent systems^36^. The ternary mixture likely represents a compromise that extracts a broad but shallower range of metabolites, whereas the optimized binary systems provide deeper coverage of the metabolite classes most abundant in each matrix.

### Clinical Application: Metabolic Signatures of Embryo Ploidy and Quality

Applying the optimized workflow to 120 clinical SECM samples revealed distinct metabolic signatures associated with both ploidy status and morphological quality. The strong discrimination between euploid and aneuploid embryos (PLS-DA clear separation; AUC = 0.977) demonstrates that the metabolic footprint of SECM carries robust information about chromosomal status. Notably, lipid metabolism pathways emerged as the dominant discriminatory signatures, with significant enrichment of α-linolenic acid metabolism, arachidonic acid metabolism, fatty acid biosynthesis and elongation, and mitochondrial β-oxidation. This is biologically plausible: early embryonic development is highly energy-intensive and dependent on fatty acid oxidation for ATP production, particularly during compaction and blastocyst formation when mitochondrial activity increases markedly^45^.

The differential abundance of specific fatty acids between euploid and aneuploid embryos—including elevated docosapentaenoic acid and adrenic acid in aneuploid samples—suggests that aneuploid embryos may experience altered lipid metabolism and membrane remodeling. Polyunsaturated fatty acids such as DHA and adrenic acid are critical for membrane fluidity, signaling, and redox homeostasis^37, 38^. Conversely, elevated saturated fatty acids such as palmitic acid and stearic acid may induce lipotoxicity and oxidative stress, impairing blastocyst formation^39^. The coordinated changes in sphingolipid metabolism and steroidogenesis further implicate apoptosis regulation and hormonal microenvironment in aneuploidy-associated metabolic dysfunction.

Amino acid and organic acid metabolism also contributed to ploidy discrimination. Methionine, a key substrate for one-carbon metabolism and epigenetic regulation via S-adenosylmethionine, was differentially expressed between groups, consistent with the known importance of epigenetic fidelity for chromosomal stability during preimplantation development^40, 41^. Similarly, alterations in TCA cycle intermediates (citric acid, isocitric acid) and energy substrates (pyruvate, lactate) reflect mitochondrial functional status, which is increasingly recognized as a critical determinant of embryo quality^44, 45^.

### Progressive Metabolic Alterations Across Morphological Grades

Analysis of euploid embryos stratified by morphological quality (good, fair, poor) revealed progressive metabolic deterioration that parallels the decline in morphological grade. Notably, the metabolic differences between good and fair embryos were primarily driven by TCA cycle activity and tryptophan/sphingolipid metabolism, whereas the differences between fair and poor embryos were dominated by lipid metabolism and membrane composition changes. This suggests a hierarchical pattern of metabolic decompensation: early-stage quality decline may manifest as mitochondrial energetic insufficiency, whereas more severe quality decline involves broader lipid metabolic dysregulation.

The enrichment of fatty acid β-oxidation, nicotinamide metabolism, and branched-chain amino acid degradation in the good vs. poor comparison indicates that low-quality embryos suffer from combined deficits in energy production, redox homeostasis, and amino acid metabolic reprogramming. The progressive involvement of these pathways as quality declines underscores the interconnected nature of metabolic networks and suggests that no single pathway is sufficient to capture embryo developmental competence; rather, a systems-level assessment is required.

### Machine Learning Models for Embryo Quality Assessment

The weighted ensemble machine learning model achieved excellent performance for both ploidy prediction (AUC = 0.977) and morphological quality classification (AUCs of 0.944, 0.889, and 0.943 for good, fair, and poor quality, respectively). The ensemble approach, which combines predictions from multiple base models weighted by their individual accuracy, outperformed any single model, demonstrating the value of leveraging complementary algorithms to capture different aspects of the metabolomics data structure.

The high specificity for euploid embryos (100.0%) is particularly noteworthy, as it suggests that the model rarely misclassifies a euploid embryo as aneuploid—a critical attribute for a clinical screening tool where false positives could lead to unnecessary discard of viable embryos. The sensitivity for aneuploidy detection (89.6%) is also encouraging, though further improvement would be desirable to minimize false negatives. These performance metrics compare favorably with or exceed those reported in previous SECM metabolomics studies using less optimized workflows^13, 16^, underscoring the importance of rigorous sample preparation optimization for achieving robust predictive models.

### Comparison with Existing Non-Invasive Approaches

Current non-invasive embryo assessment methods, including time-lapse imaging and AI-based morphological scoring, provide valuable information but remain limited in their ability to predict ploidy status and implantation potential^6–8^. PGT-A remains the gold standard for ploidy assessment but requires invasive biopsy and is associated with additional costs, procedural risks, and potential impacts on embryo viability^11, 12^. Our SECM metabolomics approach offers a non-invasive alternative that captures the functional metabolic state of the embryo rather than merely its morphological appearance. Unlike imaging-based methods, which assess static structural features, metabolomics provides dynamic information about active biochemical processes, including energy metabolism, lipid homeostasis, and oxidative stress—all of which are intimately linked to developmental competence.

The combination of metabolic profiling with machine learning represents a paradigm shift from univariate biomarker discovery to multivariate pattern recognition. Rather than relying on a single metabolite, our approach leverages the collective information from multiple metabolic features to generate a probabilistic prediction of embryo quality. This systems-level strategy is better suited to capture the complexity of embryo metabolism and is more likely to translate into clinically useful decision support tools.

### Limitations and Future Directions

Several limitations of this study should be acknowledged. First, although our sample size of 120 SECM samples is among the larger cohorts reported for SECM metabolomics, it remains modest for robust machine learning model development. External validation in independent, multi-center cohorts is essential to confirm model generalizability and to establish clinically acceptable performance thresholds. Second, our ploidy assessment was based on trophectoderm biopsy, which reflects the chromosomal status of the trophectoderm lineage but may not fully capture the inner cell mass ploidy due to mosaicism. Third, we did not prospectively evaluate implantation or live birth outcomes, which represent the ultimate clinical endpoints of interest. Future studies should incorporate clinical follow-up data to determine whether SECM metabolic signatures predict not only ploidy but also implantation success and ongoing pregnancy. Finally, the optimized protocol described here is specific to our LC-MS platform and chromatographic conditions; transfer to other platforms may require minor adjustments.

Despite these limitations, this study provides a rigorous methodological foundation for SECM metabolomics and demonstrates its clinical potential. The optimized sample preparation workflow addresses a critical gap in method standardization that has hindered cross-study comparisons and clinical translation in the field. The identification of robust metabolic signatures associated with ploidy and morphological quality, combined with the excellent performance of machine learning models, supports the feasibility of non-invasive embryo assessment using SECM metabolomics. Future large-scale, prospective, multi-center studies are warranted to validate these findings and to explore whether metabolic signatures can complement or potentially replace existing invasive approaches in select clinical scenarios. Integration of metabolomic data with morphokinetic parameters and genetic information may further improve predictive accuracy and facilitate personalized embryo selection in assisted reproductive technologies.

## Conclusions

This study systematically optimized sample preparation for untargeted LC-MS metabolomics of SECM, demonstrating that optimal extraction and reconstitution conditions are highly matrix-dependent. The optimized workflow (7× 50% ACN extraction and 40% ACN reconstitution) maximized metabolome coverage for SECM, whereas serum required 10× 100% MeOH extraction and 100% water reconstitution, highlighting that sample preparation protocols should not be directly transferred across biofluids. Application of the optimized method to 120 clinical SECM samples identified distinct metabolic signatures associated with embryonic ploidy, with lipid metabolism emerging as a key discriminatory pathway. A weighted ensemble machine-learning model achieved excellent performance for distinguishing euploid and aneuploid embryos (AUC = 0.977). Furthermore, metabolomic profiling of morphologically graded euploid embryos revealed progressive alterations in energy and lipid metabolism, with AUCs of 0.944, 0.889, and 0.943 for good-, fair-, and poor-quality embryos, respectively. Overall, this study establishes a robust workflow for SECM metabolomics and highlights its potential for non-invasive embryo ploidy screening and quality assessment in assisted reproductive technologies. Future large-scale, prospective, multi-center studies are warranted to validate these findings and assess their clinical utility.

## Supporting Information

The Supporting Information is available, free of charge at…

## Declarations

### Ethics approval and consent to participate

This study was conducted in accordance with the Declaration of Helsinki and was approved by the Ethics Committee of the Reproductive and Genetic Hospital of CITIC-XIANGYA (Approval No. LL-SC-2023-033). All participants provided written informed consent prior to sample collection.

### Consent for publication

Not applicable. This manuscript does not contain any individual person’s data in any form (including individual details, images, or videos).

### Availability of data and materials

The mass spectrometry raw files generated during this study have been deposited in the OMIX database under accession code OMIX017698 (https://bigd.big.ac.cn/omix/). The processed metabolomics data and machine learning code supporting the findings of this study are available from the corresponding author upon reasonable request.

### Authors’ contributions

S.Z., H.C., and J.Z. conceived and directed the project. S.Z. developed the concept and designed experiments. H.G., X.W., F.T., and H.I. performed most of the experiments. X.W. and H.G. analyzed the data. X.C. assisted in the metabolomics experiments. P.X., S.Z., and G.L. contributed to project conceptualization. H.G. and S.Z. wrote the manuscript. S.Z., H.C., and J.Z. revised the manuscript. All authors read and approved the final manuscript.

## Acknowledgements

The authors thank the clinical staff at the Reproductive and Genetic Hospital of CITIC-XIANGYA for their assistance with sample collection and patient management. We also thank all patients and volunteers who participated in this study.

## References

1. Steptoe, P. C.; Edwards, R. G., Birth after the reimplantation of a human embryo. Lancet 1978, 2 (8085), 366.

2. Sunderam, S.; Kissin, D. M.; Zhang, Y.; Jewett, A.; Boulet, S. L.; Warner, L.; Kroelinger, C. D.; Barfield, W. D., Assisted Reproductive Technology Surveillance - United States, 2018. MMWR Surveill Summ 2022, 71 (4), 1–19.

3. Cutting, R., Single embryo transfer for all. Best Pract Res Clin Obstet Gynaecol 2018, 53, 30–37.

4. McLernon, D. J.; Harrild, K.; Bergh, C.; Davies, M. J.; de Neubourg, D.; Dumoulin, J. C.; Gerris, J.; Kremer, J. A.; Martikainen, H.; Mol, B. W.; Norman, R. J.; Thurin-Kjellberg, A.; Tiitinen, A.; van Montfoort, A. P.; van Peperstraten, A. M.; Van Royen, E.; Bhattacharya, S., Clinical effectiveness of elective single versus double embryo transfer: meta-analysis of individual patient data from randomised trials. BMJ 2010, 341, c6945.

5. Gardner, D. K.; Lane, M.; Stevens, J.; Schlenker, T.; Schoolcraft, W. B., Blastocyst score affects implantation and pregnancy outcome: towards a single blastocyst transfer. Fertil Steril 2000, 73 (6), 1155–8.

6. Gardner, D. K.; Balaban, B., Assessment of human embryo development using morphological criteria in an era of time-lapse, algorithms and ‘OMICS’: is looking good still important? Mol Hum Reprod 2016, 22 (10), 704–718.

7. VerMilyea, M.; Hall, J. M. M.; Diakiw, S. M.; Johnston, A.; Nguyen, T.; Perugini, D.; Miller, A.; Picou, A.; Murphy, A. P.; Perugini, M., Development of an artificial intelligence-based assessment model for prediction of embryo viability using static images captured by optical light microscopy during IVF. Hum Reprod 2020, 35 (4), 770–784.

8. Liu, H.; Chen, L.; Shan, G.; Sun, C.; Lu, C.; Liao, H.; Zhang, S.; Dong, S.; Xu, X.; Yan, Q.; Gong, F.; Zhang, Z.; Dai, C.; Chen, W.; Song, H.; Chen, L.; Wang, S.; Sun, H.; Lin, G.; Sun, Y.; Gu, Y., An interpretable artificial intelligence approach to differentiate between blastocysts with similar or same morphological grades. Hum Reprod 2025, 40 (6), 1077–1086.

9. Munne, S.; Kaplan, B.; Frattarelli, J. L.; Child, T.; Nakhuda, G.; Shamma, F. N.; Silverberg, K.; Kalista, T.; Handyside, A. H.; Katz-Jaffe, M.; Wells, D.; Gordon, T.; Stock-Myer, S.; Willman, S.; Group, S. S., Preimplantation genetic testing for aneuploidy versus morphology as selection criteria for single frozen-thawed embryo transfer in good-prognosis patients: a multicenter randomized clinical trial. Fertil Steril 2019, 112 (6), 1071–1079 e7.

10. Tiegs, A. W.; Tao, X.; Zhan, Y.; Whitehead, C.; Kim, J.; Hanson, B.; Osman, E.; Kim, T. J.; Patounakis, G.; Gutmann, J.; Castelbaum, A.; Seli, E.; Jalas, C.; Scott, R. T., Jr., A multicenter, prospective, blinded, nonselection study evaluating the predictive value of an aneuploid diagnosis using a targeted next-generation sequencing-based preimplantation genetic testing for aneuploidy assay and impact of biopsy. Fertil Steril 2021, 115 (3), 627–637.

11. Scott, R. T., Jr.; Upham, K. M.; Forman, E. J.; Hong, K. H.; Scott, K. L.; Taylor, D.; Tao, X.; Treff, N. R., Blastocyst biopsy with comprehensive chromosome screening and fresh embryo transfer significantly increases in vitro fertilization implantation and delivery rates: a randomized controlled trial. Fertil Steril 2013, 100 (3), 697–703.

12. Zhang, S.; Luo, K.; Cheng, D.; Tan, Y.; Lu, C.; He, H.; Gu, Y.; Lu, G.; Gong, F.; Lin, G., Number of biopsied trophectoderm cells is likely to affect the implantation potential of blastocysts with poor trophectoderm quality. Fertil Steril 2016, 105 (5), 1222–1227 e4.

13. Zmuidinaite, R.; Sharara, F. I.; Iles, R. K., Current Advancements in Noninvasive Profiling of the Embryo Culture Media Secretome. Int J Mol Sci 2021, 22 (5).

14. Bracewell-Milnes, T.; Saso, S.; Abdalla, H.; Nikolau, D.; Norman-Taylor, J.; Johnson, M.; Holmes, E.; Thum, M. Y., Metabolomics as a tool to identify biomarkers to predict and improve outcomes in reproductive medicine: a systematic review. Hum Reprod Update 2017, 23 (6), 723–736.

15. Leese, H. J.; Brison, D. R.; Sturmey, R. G., The Quiet Embryo Hypothesis: 20 years on. Front Physiol 2022, 13, 899485.

16. Ferrick, L.; Lee, Y. S. L.; Gardner, D. K., Metabolic activity of human blastocysts correlates with their morphokinetics, morphological grade, KIDScore and artificial intelligence ranking. Hum Reprod 2020, 35 (9), 2004–2016.

17. Gardner, D. K.; Lane, M.; Stevens, J.; Schoolcraft, W. B., Noninvasive assessment of human embryo nutrient consumption as a measure of developmental potential. Fertil Steril 2001, 76 (6), 1175–80.

18. Picton, H. M.; Elder, K.; Houghton, F. D.; Hawkhead, J. A.; Rutherford, A. J.; Hogg, J. E.; Leese, H. J.; Harris, S. E., Association between amino acid turnover and chromosome aneuploidy during human preimplantation embryo development in vitro. Mol Hum Reprod 2010, 16 (8), 557–69.

19. Wallace, M.; Cottell, E.; Cullinane, J.; McAuliffe, F. M.; Wingfield, M.; Brennan, L., (1)H NMR based metabolic profiling of day 2 spent embryo media correlates with implantation potential. Syst Biol Reprod Med 2014, 60 (1), 58–63.

20. Patti, G. J.; Yanes, O.; Siuzdak, G., Innovation: Metabolomics: the apogee of the omics trilogy. Nat Rev Mol Cell Biol 2012, 13 (4), 263–9.

21. Giera, M.; Yanes, O.; Siuzdak, G., Metabolite discovery: Biochemistry’s scientific driver. Cell Metab 2022, 34 (1), 21–34.

22. Dyrlund, T. F.; Kirkegaard, K.; Poulsen, E. T.; Sanggaard, K. W.; Hindkjaer, J. J.; Kjems, J.; Enghild, J. J.; Ingerslev, H. J., Unconditioned commercial embryo culture media contain a large variety of non-declared proteins: a comprehensive proteomics analysis. Hum Reprod 2014, 29 (11), 2421–30.

23. Morbeck, D. E.; Krisher, R. L.; Herrick, J. R.; Baumann, N. A.; Matern, D.; Moyer, T., Composition of commercial media used for human embryo culture. Fertil Steril 2014, 102 (3), 759–766 e9.

24. Dunn, W. B.; Broadhurst, D.; Begley, P.; Zelena, E.; Francis-McIntyre, S.; Anderson, N.; Brown, M.; Knowles, J. D.; Halsall, A.; Haselden, J. N.; Nicholls, A. W.; Wilson, I. D.; Kell, D. B.; Goodacre, R.; Human Serum Metabolome, C., Procedures for large-scale metabolic profiling of serum and plasma using gas chromatography and liquid chromatography coupled to mass spectrometry. Nat Protoc 2011, 6 (7), 1060–83.

25. Want, E. J.; Masson, P.; Michopoulos, F.; Wilson, I. D.; Theodoridis, G.; Plumb, R. S.; Shockcor, J.; Loftus, N.; Holmes, E.; Nicholson, J. K., Global metabolic profiling of animal and human tissues via UPLC-MS. Nat Protoc 2013, 8 (1), 17–32.

26. Vuckovic, D., Current trends and challenges in sample preparation for global metabolomics using liquid chromatography-mass spectrometry. Anal Bioanal Chem 2012, 403 (6), 1523–48.

27. Bruce, S. J.; Tavazzi, I.; Parisod, V.; Rezzi, S.; Kochhar, S.; Guy, P. A., Investigation of human blood plasma sample preparation for performing metabolomics using ultrahigh performance liquid chromatography/mass spectrometry. Anal Chem 2009, 81 (9), 3285–96.

28. Tulipani, S.; Llorach, R.; Urpi-Sarda, M.; Andres-Lacueva, C., Comparative analysis of sample preparation methods to handle the complexity of the blood fluid metabolome: when less is more. Anal Chem 2013, 85 (1), 341–8.

29. Yanes, O.; Tautenhahn, R.; Patti, G. J.; Siuzdak, G., Expanding coverage of the metabolome for global metabolite profiling. Anal Chem 2011, 83 (6), 2152–61.

30. Whiley, L.; Godzien, J.; Ruperez, F. J.; Legido-Quigley, C.; Barbas, C., In-vial dual extraction for direct LC-MS analysis of plasma for comprehensive and highly reproducible metabolic fingerprinting. Anal Chem 2012, 84 (14), 5992–9.

31. Masson, P.; Alves, A. C.; Ebbels, T. M.; Nicholson, J. K.; Want, E. J., Optimization and evaluation of metabolite extraction protocols for untargeted metabolic profiling of liver samples by UPLC-MS. Anal Chem 2010, 82 (18), 7779–86.

32. Garcia-Canaveras, J. C.; Lopez, S.; Castell, J. V.; Donato, M. T.; Lahoz, A., Extending metabolome coverage for untargeted metabolite profiling of adherent cultured hepatic cells. Anal Bioanal Chem 2016, 408 (4), 1217–30.

33. Lindahl, A.; Saaf, S.; Lehtio, J.; Nordstrom, A., Tuning Metabolome Coverage in Reversed Phase LC-MS Metabolomics of MeOH Extracted Samples Using the Reconstitution Solvent Composition. Anal Chem 2017, 89 (14), 7356–7364.

34. Siristatidis, C.; Dafopoulos, K.; Papapanou, M.; Stavros, S.; Pouliakis, A.; Eleftheriades, A.; Sidiropoulou, T.; Vlahos, N., Why Has Metabolomics So Far Not Managed to Efficiently Contribute to the Improvement of Assisted Reproduction Outcomes? The Answer through a Review of the Best Available Current Evidence. Diagnostics (Basel) 2021, 11 (9).

35. Fluss, R.; Faraggi, D.; Reiser, B., Estimation of the Youden Index and its associated cutoff point. Biom J 2005, 47 (4), 458–72.

36. Dettmer, K.; Nurnberger, N.; Kaspar, H.; Gruber, M. A.; Almstetter, M. F.; Oefner, P. J., Metabolite extraction from adherently growing mammalian cells for metabolomics studies: optimization of harvesting and extraction protocols. Anal Bioanal Chem 2011, 399 (3), 1127–39.

37. Yagi, A.; Miyanaga, S.; Shrestha, R.; Takeda, S.; Kobayashi, S.; Chiba, H.; Kamiya, H.; Hui, S. P., A fatty acid profiling method using liquid chromatography-high resolution mass spectrometry for improvement of assisted reproductive technology. Clin Chim Acta 2016, 456, 100–106.

38. Crawford, M. A.; Sinclair, A. J.; Hall, B.; Ogundipe, E.; Wang, Y.; Bitsanis, D.; Djahanbakhch, O. B.; Harbige, L.; Ghebremeskel, K.; Golfetto, I.; Moodley, T.; Hassam, A.; Sassine, A.; Johnson, M. R., The imperative of arachidonic acid in early human development. Prog Lipid Res 2023, 91, 101222.

39. Saeed, H. A.; Sabir, R.; Lu, X.; Jiang, Y.; Koutonin, B. O. M.; Wang, D.; Fu, Y.; Jia, C.; Li, J., 6-Gingerol and Astaxanthin Mitigate the Effects of Stearic Acid in Pig Oocyte Maturation. Reprod Domest Anim 2024, 59 (11), e14746.

40. Sagheer, M.; Carballo, D.; Maia, T. S.; Hansen, P. J., Consequences of varying methionine concentrations on development of the bovine embryo in vitrodagger. Biol Reprod 2025, 113 (4), 765–776.

41. Saha, S.; Debacq, C.; Audouard, C.; Jungas, T.; Dupre, P.; Mohamad-Ali, F.; Chapat, C.; Michaud, H. A.; Le Cam, L.; Lacroix, M.; Ohayon, D.; Davy, A., Acute dietary methionine restriction triggers cell cycle arrest and reversible growth defects in the neocortex. iScience 2025, 28 (6), 112705.

42. Robinson, J. J., Energy requirements of ewes during late pregnancy and early lactation. Vet Rec 1980, 106 (13), 282–4.

43. Thompson, J. G.; Bell, A. C.; Pugh, P. A.; Tervit, H. R., Metabolism of pyruvate by pre-elongation sheep embryos and effect of pyruvate and lactate concentrations during culture in vitro. Reprod Fertil Dev 1993, 5 (4), 417–23.

44. Pendleton, A. L.; Antolic, A. T.; Kelly, A. C.; Davis, M. A.; Camacho, L. E.; Doubleday, K.; Anderson, M. J.; Langlais, P. R.; Lynch, R. M.; Limesand, S. W., Lower oxygen consumption and Complex I activity in mitochondria isolated from skeletal muscle of fetal sheep with intrauterine growth restriction. Am J Physiol Endocrinol Metab 2020, 319 (1), E67–E80.

45. Wakefield, S. L.; Lane, M.; Mitchell, M., Impaired mitochondrial function in the preimplantation embryo perturbs fetal and placental development in the mouse. Biol Reprod 2011, 84 (3), 572–80.

46. Dunning, K. R.; Russell, D. L.; Robker, R. L., Lipids and oocyte developmental competence: the role of fatty acids and beta-oxidation. Reproduction 2014, 148 (1), R15–27.

47. May-Panloup, P.; Boucret, L.; Chao de la Barca, J. M.; Desquiret-Dumas, V.; Ferre-L’Hotellier, V.; Moriniere, C.; Descamps, P.; Procaccio, V.; Reynier, P., Ovarian ageing: the role of mitochondria in oocytes and follicles. Hum Reprod Update 2016, 22 (6), 725–743.

48. Zhao, J.; Wang, W.; Zhang, L.; Zhang, J.; Sturmey, R.; Zhang, J., Dynamic metabolism during early mammalian embryogenesis. Development 2023, 150 (20).

49. Liu, A.; Shen, H.; Li, Q.; He, J.; Wang, B.; Du, W.; Li, G.; Zhang, M.; Zhang, X., Determination of tryptophan and its indole metabolites in follicular fluid of women with diminished ovarian reserve. Sci Rep 2023, 13 (1), 17124.

50. Kalo, D.; Roth, Z., Involvement of the sphingolipid ceramide in heat-shock-induced apoptosis of bovine oocytes. Reprod Fertil Dev 2011, 23 (7), 876–88.

51. Li, J.; Zhang, J.; Hou, W.; Yang, X.; Liu, X.; Zhang, Y.; Gao, M.; Zong, M.; Dong, Z.; Liu, Z.; Shen, J.; Cong, W.; Ding, C.; Gao, S.; Huang, G.; Kong, Q., Metabolic control of histone acetylation for precise and timely regulation of minor ZGA in early mammalian embryos. Cell Discov 2022, 8 (1), 96.

52. Xu, Y.; Xie, W.; Zhang, J., Metabolic regulation of key developmental events during mammalian embryogenesis. Nat Cell Biol 2025, 27 (8), 1219–1229.

53. Fu, J.; Gaetani, S.; Oveisi, F.; Lo Verme, J.; Serrano, A.; Rodriguez De Fonseca, F.; Rosengarth, A.; Luecke, H.; Di Giacomo, B.; Tarzia, G.; Piomelli, D., Oleylethanolamide regulates feeding and body weight through activation of the nuclear receptor PPAR-alpha. Nature 2003, 425 (6953), 90–3.

54. McKeegan, P. J.; Sturmey, R. G., The role of fatty acids in oocyte and early embryo development. Reprod Fertil Dev 2011, 24 (1), 59–67.

55. Agarwal, A.; Gupta, S.; Sharma, R. K., Role of oxidative stress in female reproduction. Reprod Biol Endocrinol 2005, 3, 28.

56. Wu, G.; Bazer, F. W.; Datta, S.; Johnson, G. A.; Li, P.; Satterfield, M. C.; Spencer, T. E., Proline metabolism in the conceptus: implications for fetal growth and development. Amino Acids 2008, 35 (4), 691–702.

57. Tsafriri, A.; Chun, S. Y.; Zhang, R.; Hsueh, A. J.; Conti, M., Oocyte maturation involves compartmentalization and opposing changes of cAMP levels in follicular somatic and germ cells: studies using selective phosphodiesterase inhibitors. Dev Biol 1996, 178 (2), 393–402.

58. Richard, F. J.; Tsafriri, A.; Conti, M., Role of phosphodiesterase type 3A in rat oocyte maturation. Biol Reprod 2001, 65 (5), 1444–51.

