## Supplemental Figure 1-11; Supplemental Table 1-8 for "Non-Invasive Embryo Quality Assessment via Matrix-Optimized Untargeted LC-MS Metabolomics of Spent Embryo Culture Media and Weighted Ensemble Machine Learning"

### Table of Content


| Description | Page No. |
|---|---|
| **Experimental section** | S-4 |
| **Figure S1.** The coefficient of variation (CV) distribution of peak area across replicates by different extraction solvent composition and volume for SECM | S-6 |
| **Figure S2.** Relative recovery of different metabolite class in SECM obtained with different extraction solvent composition and volume using 50% ACN with 7× volume as a reference | S-7 |
| **Figure S3.** The coefficient of variation (CV) distribution of peak area across replicates by different extraction solvent composition and volume for serum | S-8 |
| **Figure S4.** Relative recovery of different metabolite class in serum obtained with different extraction solvent composition and volume using 100% MeOH with 10× volume as a reference | S-9 |
| **Figure S5.** Comparison of the binary and ternary extraction solvent systems for SECM and serum | S-10 |
| **Figure S6.** Relative recovery of different metabolite class in SECM obtained with different reconsitution solvent composition using 40% ACN as a reference | S-11 |
| **Figure S7.** Relative recovery of different metabolite class in serum obtained with different reconsitution solvent composition using 0% MeOH as a reference | S-12 |
| **Figure S8.** Comparison of the profile and centroid MS acquisition modes for metabolite analysis | S-13 |
| **Figure S9.** The relative abundance differences of fifteen candidate biomarkers screened based on univariate and multivariate statistical criteria and analytical stability in E and A groups ($\|FC\| > 1.2$, $p < 0.05$,$CV< 0.25$) | S-14 |
| **Figure S10.** Heatmaps of relative concentrations (Z-score) and fold changes for identified metabolites (fatty acids, amino acids, organic acids) | S-15 |
| **Figure S11.** The relative abundance differences of twelve candidate biomarkers screened based on univariate and multivariate statistical criteria and analytical stability in G, F, and P groups ($\|FC\| > 1.2$, $p < 0.05$,$CV< 0.15$) | S-16 |

| | |
|---|---|
| **Table S1.** The number of identified metabolites from SECM by different extraction solvent composition and volume | S-17 |
| **Table S2.** The number of identified metabolites from serum by different extraction solvent composition and volume | S-18 |
| **Table S3.** The detailed information of the blastocysts corresponding to 120 SECM samples | S-19 |
| **Table S4.** The fifteen candidate biomarkers preselected based on univariate and multivariate statistical criteria and analytical stability ($\|FC\| > 1.2$, $p < 0.05$, $CV < 0.25$) | S-22 |
| **Table S5.** Performance metrics for each model, including accuracy, sensitivity, specificity, PPV, NPV, AUC, and F1 score | S-23 |
| **Table S6.** Twelve candidate biomarkers preselected based on univariate and multivariate statistical criteria and analytical stability ($\|FC\| > 1.2$, $p < 0.05$, $CV < 0.15$) | S-24 |
| **Table S7.** Performance metrics for each model, including overall accuracy, Kappa coefficient, macro-average sensitivity, specificity, PPV, NPV, F1 score, and AUC | S-25 |
| **Table S8.** Class-specific performance metrics, including sensitivity, specificity, PPV, NPV, F1 score and AUC | S-26 |
| **Reference** | S-27 |

### Methods

#### Chemicals and Reagents

LC-MS grade methanol, acetonitrile, water, and formic acid were purchased from Thermo Fisher Scientific (Waltham, MA, USA). Ammonium formate was obtained from Sigma-Aldrich (St. Louis, MO, USA). Ultrapure water was generated using a Milli-Q Integral Water Purification System (Merck Millipore, Burlington, MA, USA).

#### LC-MS/MS Analysis

Non-targeted metabolomics analysis was performed using a Vanquish Flex UHPLC system coupled to an Orbitrap Exploris 120 mass spectrometer (Thermo Fisher Scientific, Bremen, Germany). Chromatographic separation was achieved on a Hypersil GOLD™ VANQUISH™ C18 UHPLC column (100 × 2.1 mm, 1.9 µm particle size; Thermo Fisher Scientific) maintained at 40°C. The autosampler temperature was set to 4°C. The mobile phase consisted of (A) water containing 5 mM ammonium formate and 0.05% formic acid, and (B) ACN. The gradient elution program was as follows: 0–2 min, 2% B; 2–7 min, linear increase to 70% B; 7–14 min, linear increase to 90% B; 14–16 min, linear increase to 100% B; 16–20 min, hold at 100% B; 20–20.1 min, return to 2% B; 20.1–25 min, re-equilibration at 2% B. The flow rate was 0.3 mL/min.

The mass spectrometer was operated in both positive and negative electrospray ionization (ESI) modes with the following parameters: spray voltage, 3500 V (+ESI) and 2500 V (−ESI); sheath gas flow rate, 50 arbitrary units; auxiliary gas flow rate, 15 arbitrary units; sweep gas flow rate, 1 arbitrary unit; ion transfer tube temperature, 325°C; vaporizer temperature, 400°C. Full-scan MS spectra were acquired at a resolution of 60,000 (FWHM at m/z 200) over a scan range of m/z 80–1000. Data-dependent MS/MS (ddMS2) acquisition was performed using a Top N method (N=4) with normalized collision energies (NCE) of 15, 30, and 45 eV. MS/MS spectra were acquired at a resolution of 15,000. For comparison of acquisition modes, parallel analyses were performed using both profile and centroid data acquisition. Quality control (QC) samples, prepared by pooling equal aliquots from all samples within each experimental batch, were injected at regular intervals (every 12 injections) throughout the analytical run to monitor system stability and performance.

#### Data Processing and Metabolite Identification

Raw LC-MS data were processed using Compound Discoverer 3.3 software (Thermo Fisher Scientific). The processing workflow included retention time alignment, peak detection, compound grouping, and gap filling. Metabolite identification was performed by matching accurate mass (mass tolerance ±10 ppm), retention time (RT tolerance ±0.25 min), and MS/MS fragmentation patterns against multiple databases including mzCloud (https://www.mzcloud.org), mzVault, and ChemSpider. For mzCloud matching, spectral similarity scores >80 were required for positive identification. Peak areas were extracted for all detected features using a mass tolerance of ±10 ppm and retention time tolerance of ±0.25 min. Data were filtered to remove features present in less than 50% of QC samples or with coefficient of variation (CV) >30% in QC samples. For quantitative comparisons, peak areas were normalized to total ion current. Principal component analysis (PCA), partial least squares discriminant analysis (PLS-DA), and loading plots were performed using RStudio (version 4.5.1). Differential metabolites were selected based on |fold change (FC)| > 1.2 and $p < 0.05$. Pathway enrichment analysis of characteristic differential metabolites was conducted using MetaboAnalyst (https://www.metaboanalyst.ca).

**Statistical Analysis**

All statistical analyses were performed using R software (version4.5.1; R Foundation for Statistical Computing, Vienna, Austria) and GraphPad Prism (version 9.0; GraphPad Software, San Diego, CA, USA). Data are presented as mean ± standard deviation (SD) unless otherwise indicated. For comparison of metabolite identification numbers across multiple conditions, one-way ANOVA followed by Tukey's post-hoc test was applied. For pairwise comparisons, Student's t-test was used. Reproducibility was assessed by calculating the coefficient of variation (CV = SD/mean) for metabolite peak areas across technical replicates. Principal component analysis (PCA) was performed on Pareto-scaled data to visualize overall metabolic differences between extraction conditions. For relative recovery calculations, metabolite intensities from each method were normalized to those obtained using the reference method. Metabolite classification and logP values were obtained from the Human Metabolome Database (HMDB; https://hmdb.ca)[1] and PubChem (https://pubchem.ncbi.nlm.nih.gov)[2]. Pathway enrichment analysis was performed using MetaboAnalyst 5.0[3]. Statistical significance was set at $p < 0.05$.

**Figure S1. The coefficient of variation (CV) distribution of peak area across replicates by different extraction solvent composition and volume for SECM.**

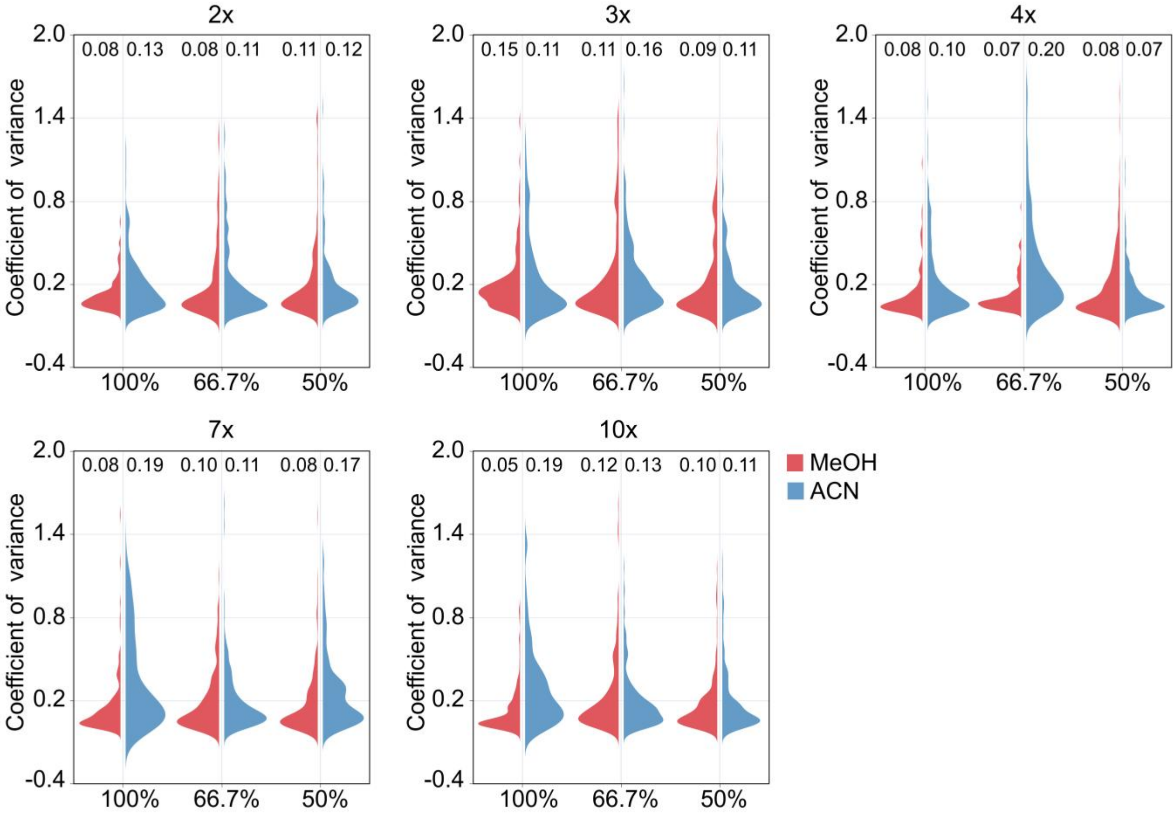

**Figure S2. Relative recovery of different metabolite class in SECM obtained with different extraction solvent composition and volume using 50% ACN with 7× volume as a reference.** (A) Relative recovery of amino acids, peptides and analogues. (B) Relative recovery of benzenoids. (C) Relative recovery of fatty acyls. (D) Relative recovery of indoles and derivatives. (E) Relative recovery of organonitrogen compounds. (F) Relative recovery of organooxygen compounds.

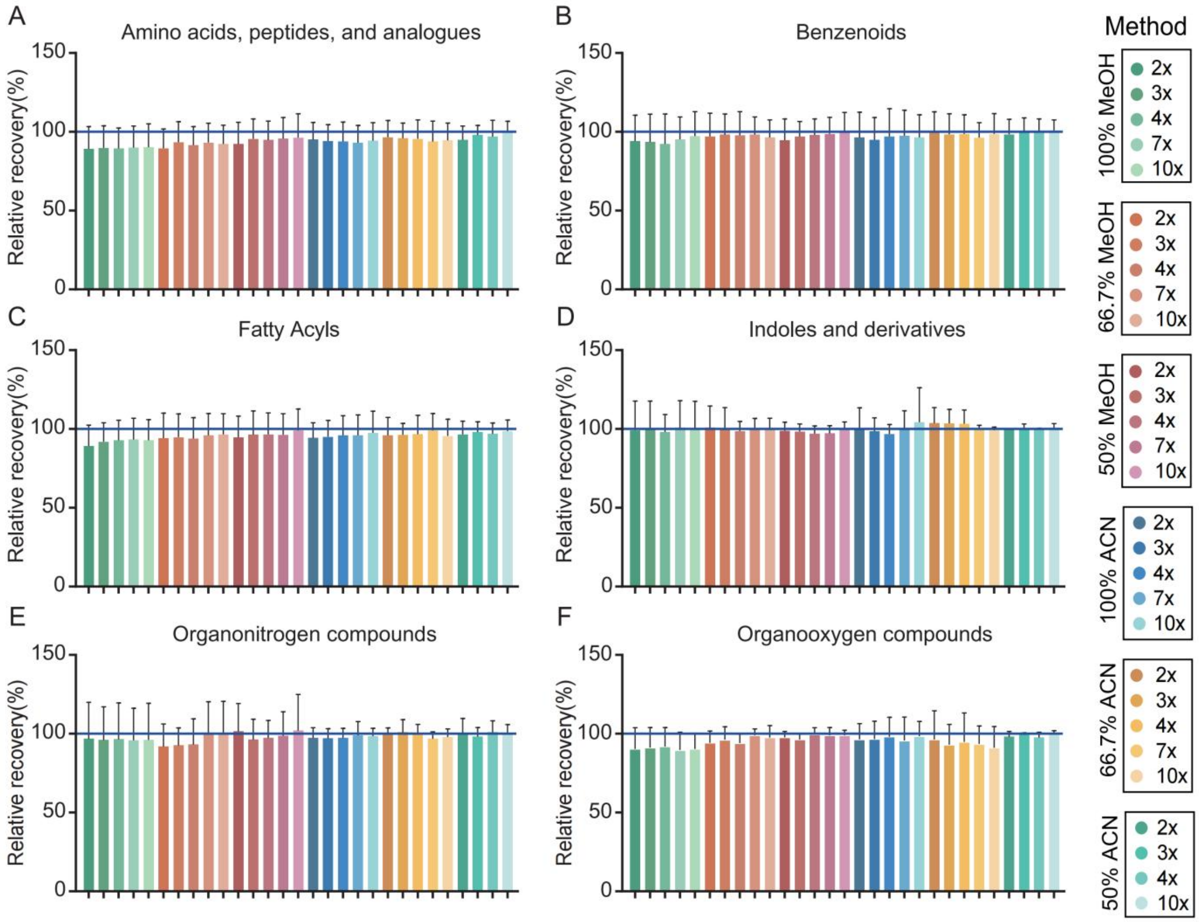

**Figure S3. The coefficient of variation (CV) distribution of peak area across replicates by different extraction solvent composition and volume for serum.**

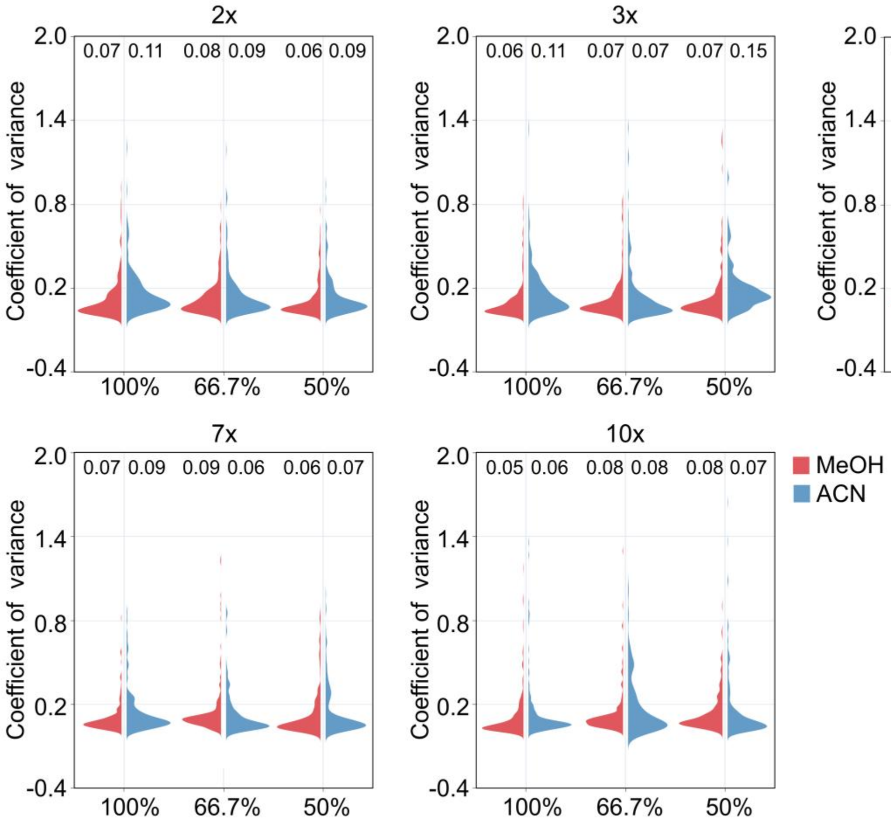

**Figure S4. Relative recovery of different metabolite class in serum obtained with different extraction solvent composition and volume using 100% MeOH with 10× volume as a reference.** (A) Relative recovery of amino acids, peptides and analogues. (B) Relative recovery of benzenoids. (C) Relative recovery of fatty acyls. (D) Relative recovery of hydroxy acids and derivatives. (E) Relative recovery of indoles and derivatives. (F) Relative recovery of organooxygen compounds.

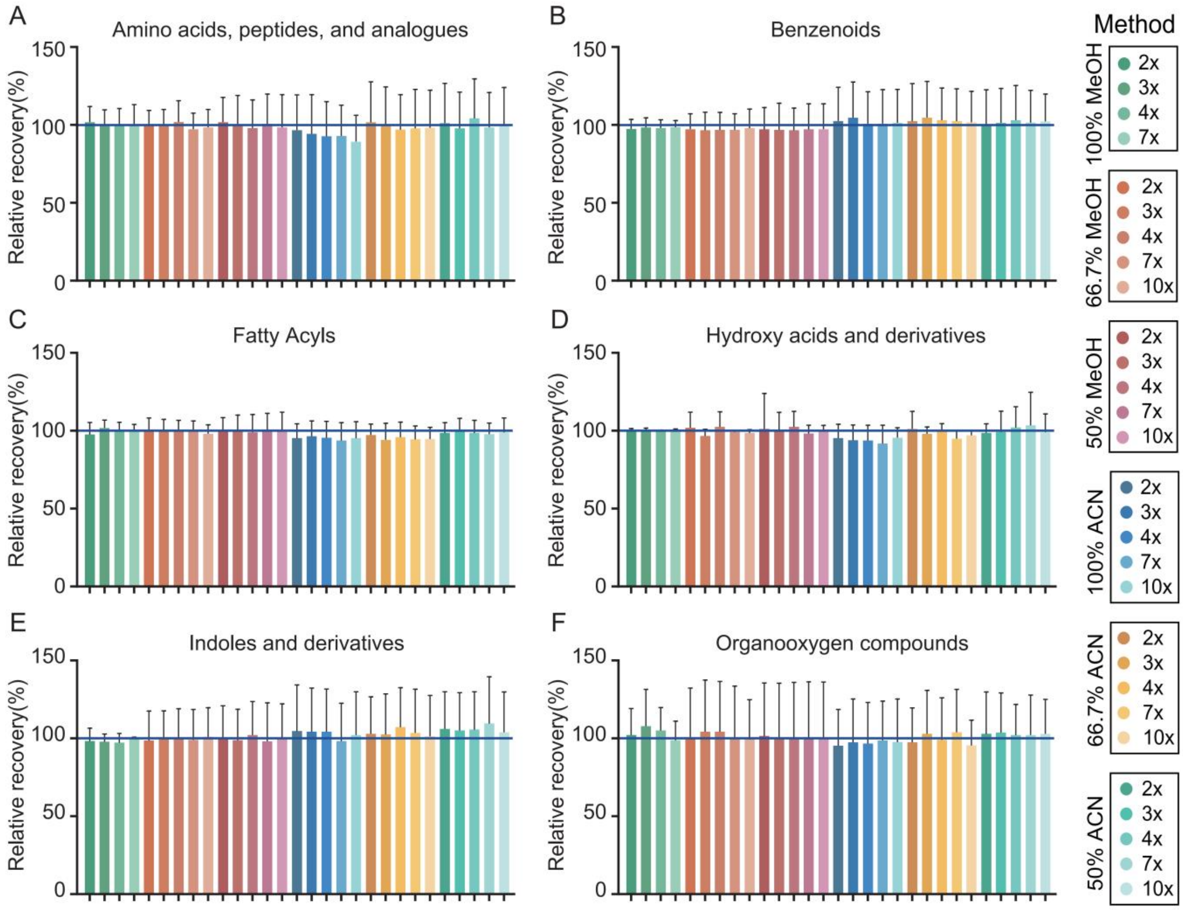

**Figure S5. Comparison of the binary and ternary extraction solvent systems for SECM and serum.** (A) The metabolite identification numbers of the optimized binary solvent system and two ternary solvent systems for SECM. Error bars: standard error (n = 3). Statistical significance was assessed by one-way ANOVA (**p < 0.01, ***p < 0.001, ****p < 0.0001 and n.s > 0.05). (B) The overlap of the identified metabolites by the optimized binary solvent system and two ternary solvent systems for SECM. (C) The coefficient of variation (CV) distribution of peak area across replicates by the optimized binary solvent system and two ternary solvent systems for SECM. (D) The metabolite identification numbers of the optimized binary solvent system and two ternary solvent systems for serum. (E) The overlap of the identified metabolites by the optimized binary solvent system and two ternary solvent systems for serum. (F) The coefficient of variation (CV) distribution of peak area across replicates by the optimized binary solvent system and two ternary solvent systems for serum.

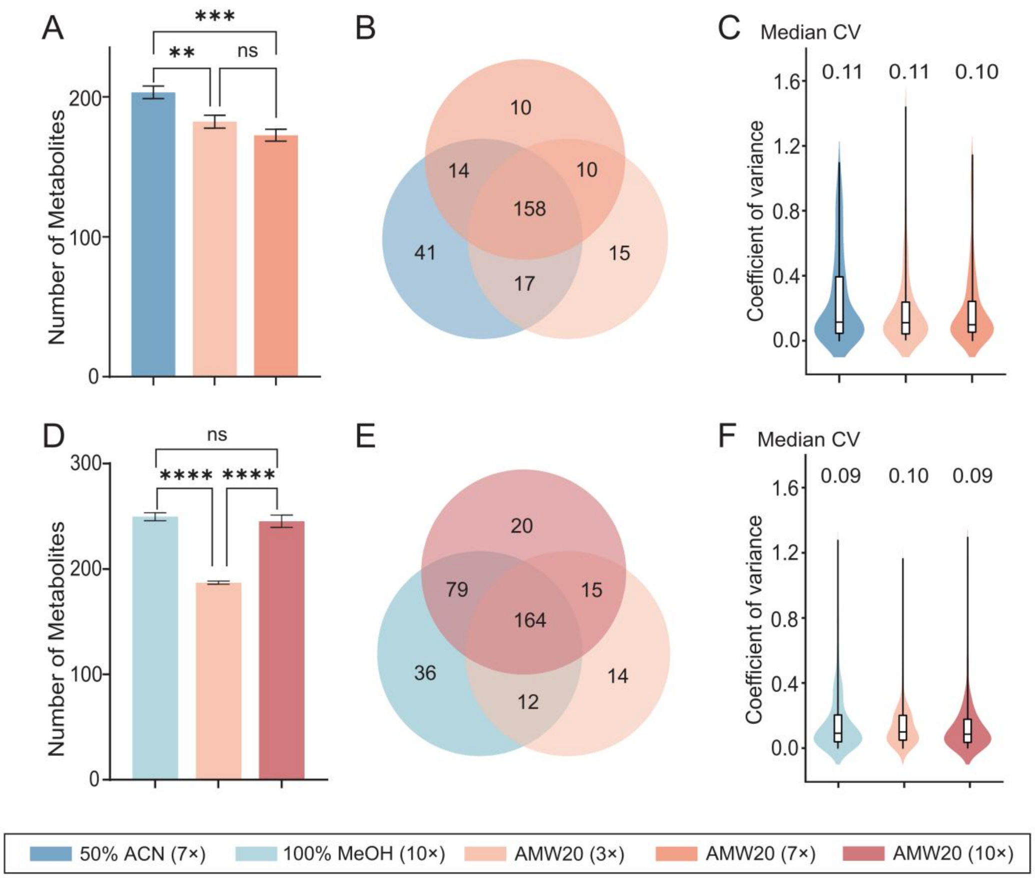

**Figure S6. Relative recovery of different metabolite class in SECM obtained with different reconsitution solvent composition using 40% ACN as a reference.** (A) Relative recovery of amino acids, peptides and analogues. (B) Relative recovery of benzenoids. (C) Relative recovery of fatty acyls. (D) Relative recovery of hydroxy acids and derivatives. (E) Relative recovery of organic phosphoric acids and derivatives. (F) Relative recovery of steroids and steroid derivatives.

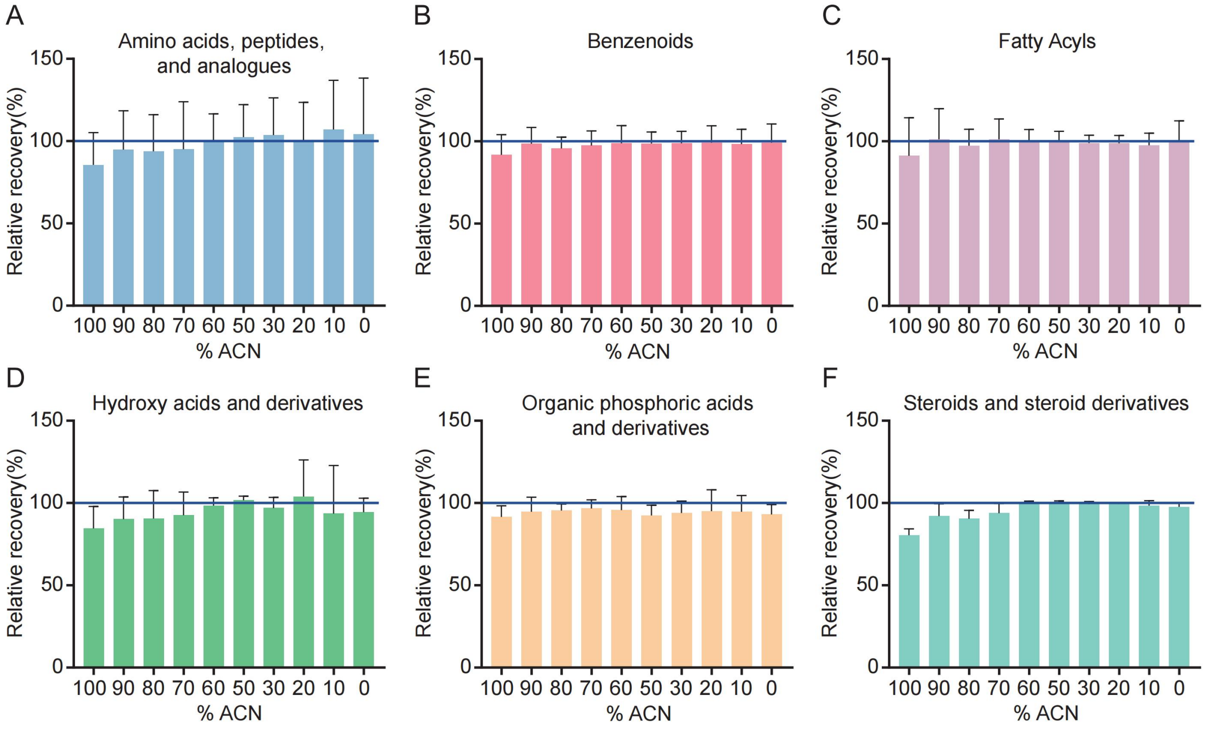

**Figure S7. Relative recovery of different metabolite class in serum obtained with different reconsitution solvent composition using 0% MeOH as a reference.** (A) Relative recovery of amino acids, peptides and analogues. (B) Relative recovery of benzenoids. (C) Relative recovery of fatty acyls. (D) Relative recovery of hydroxy acids and derivatives. (E) Relative recovery of indoles and derivatives. (F) Relative recovery of imidazopyrimidines.

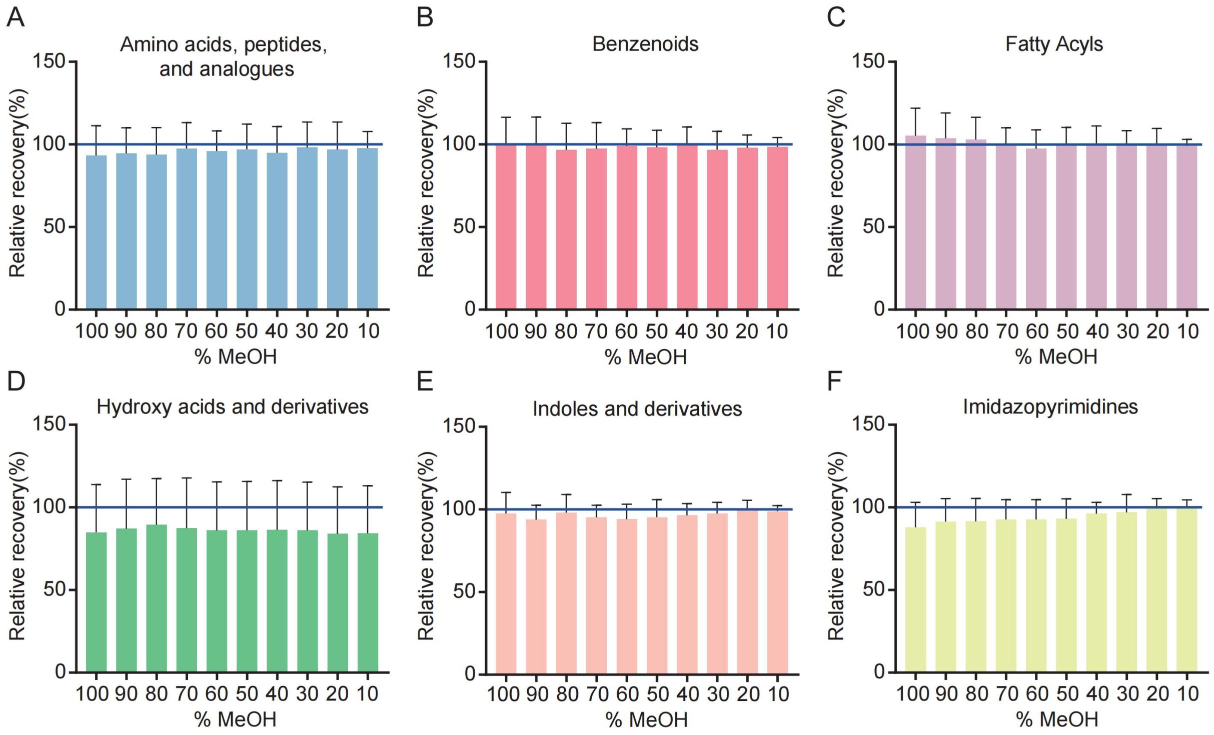

**Figure S8. Comparison of the profile and centroid MS acquisition modes for metabolite analysis.** (A) The metabolite identification numbers by profile and centroid MS acquisition modes with two reconstitution solvents. Error bars: standard error (n = 3). Statistical significance was assessed by two-way ANOVA (**p < 0.01). (B) The overlap of the identified metabolites by profile and centroid MS acquisition modes with two reconstitution solvents. (C) The coefficient of variation (CV) distribution of peak area across replicates by profile and centroid MS acquisition modes with two reconstitution solvents.

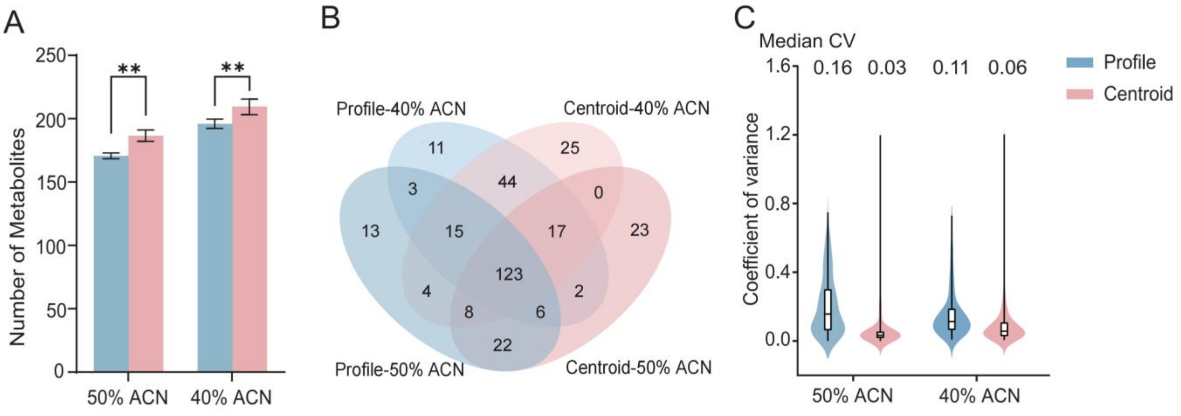

**Figure S9. The relative abundance differences of fifteen candidate biomarkers screened based on univariate and multivariate statistical criteria and analytical stability in E and A groups (|FC| > 1.2, p < 0.05,CV< 0.25). *p < 0.05, **p < 0.01, ***p < 0.001 and ****p < 0.0001 (Student's t-test).**

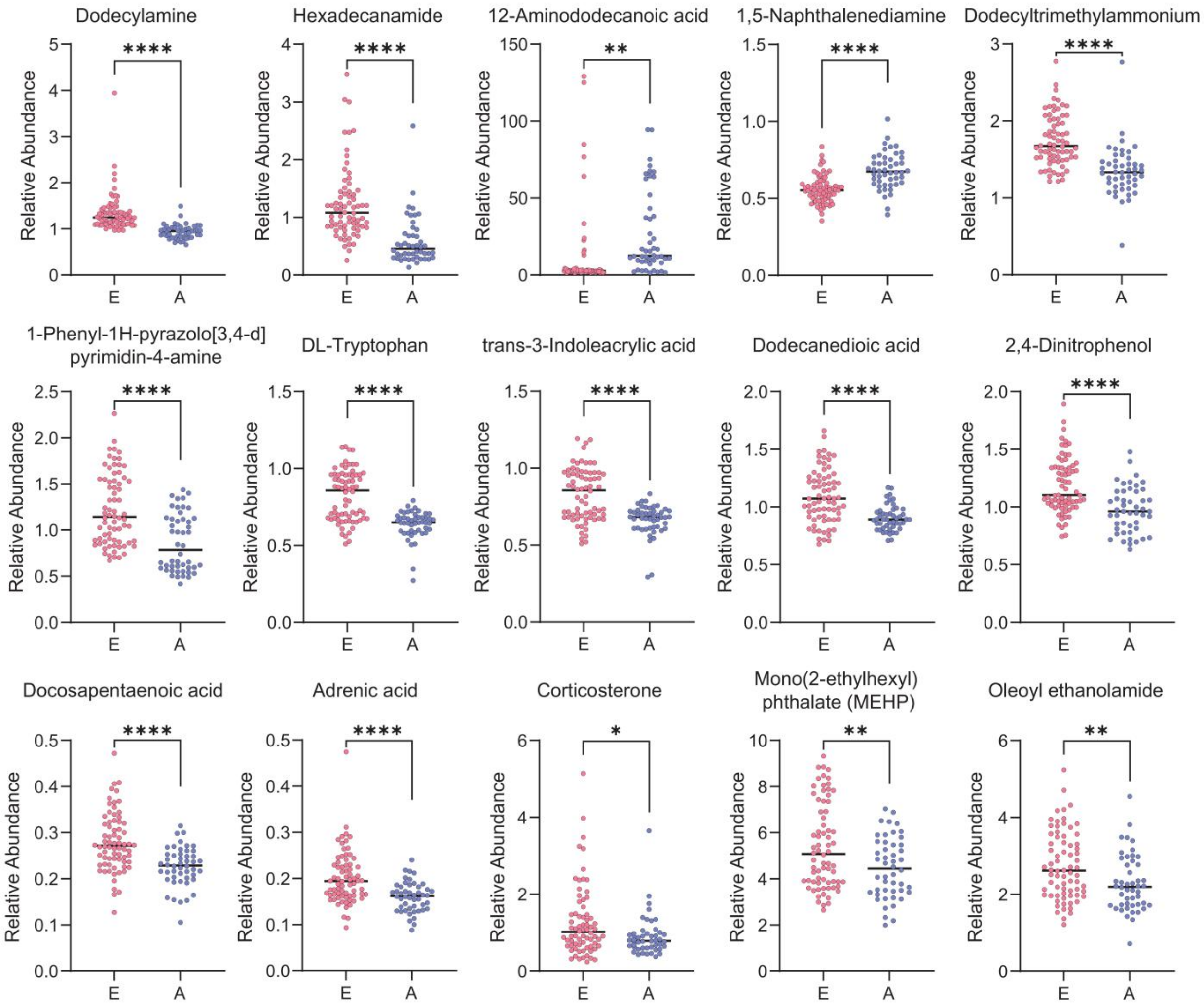

**Figure S10. Heatmaps of relative concentrations (Z-score) and fold changes for identified metabolites (fatty acids, amino acids, organic acids).** *p < 0.05, **p < 0.01, ***p < 0.001 (Student's t-test). Z-score normalization was applied to better display the relative concentrations of the mean values of each metabolite among the three groups.

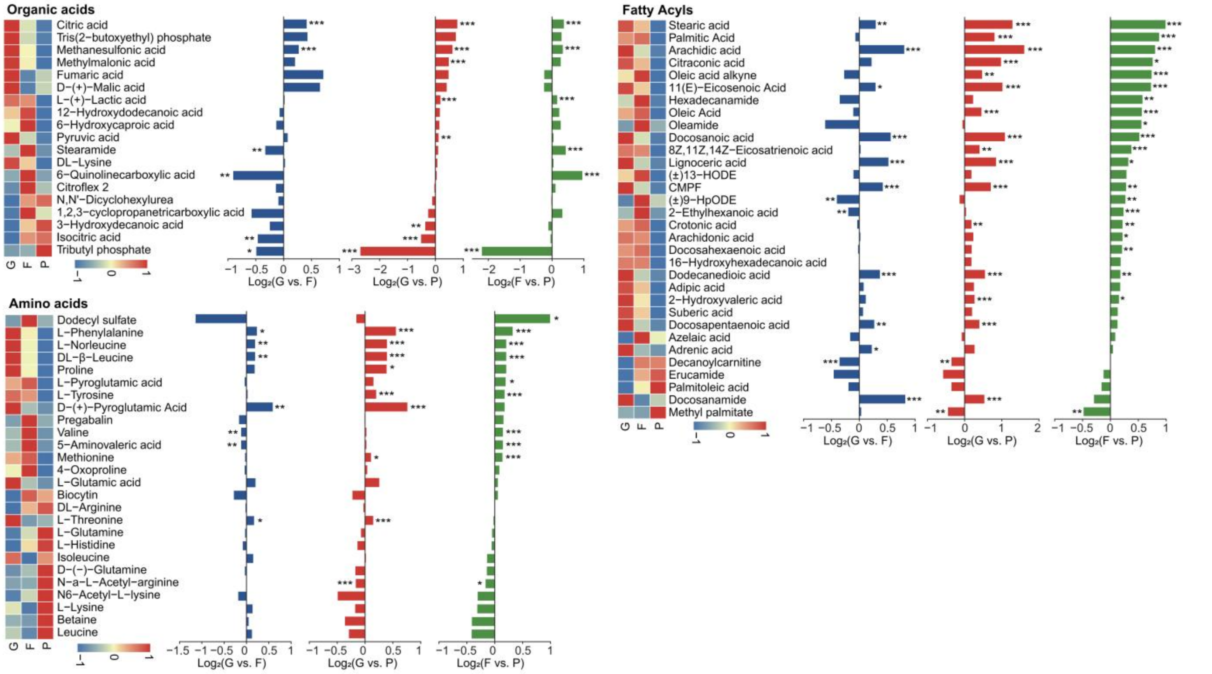

**Figure S11. The relative abundance differences of twelve candidate biomarkers screened based on univariate and multivariate statistical criteria and analytical stability in G, F, and P groups (|FC| > 1.2, p < 0.05,CV< 0.15). *p < 0.05, **p < 0.01, ***p < 0.001 and ****p < 0.0001 (Student's t-test).**

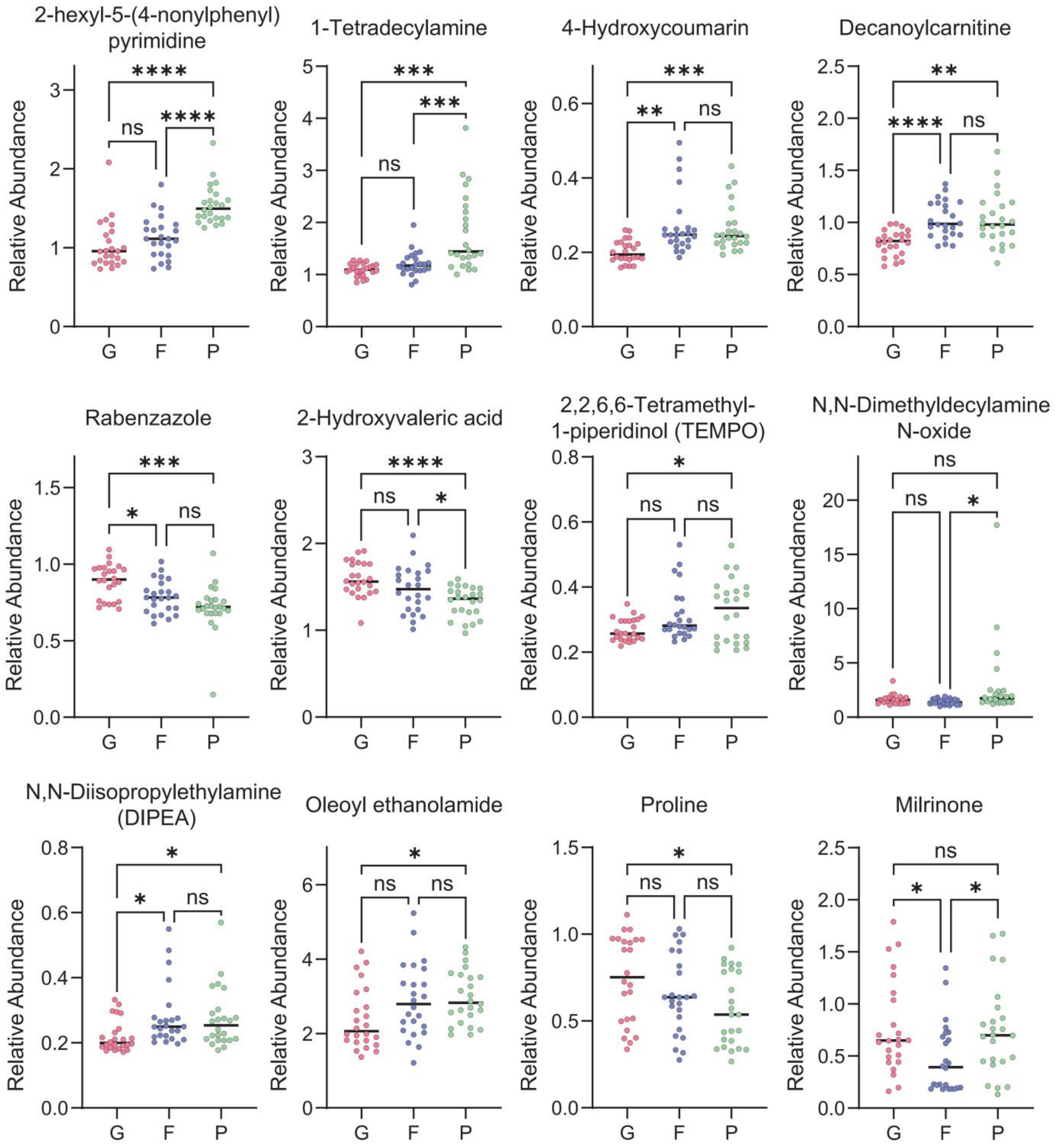

**Table S1. The number of identified metabolites from SECM by different extraction solvent composition and volume.**

| Protein precipitation solvent composition | Extraction solvent volume | Identification number |
|---|---|---|
| 100% MeOH | 2× | 134±1 |
| | 3× | 145±4 |
| | 4× | 146±4 |
| | 7× | 138±0 |
| | 10× | 138±2 |
| 66.7% MeOH | 2× | 138±5 |
| | 3× | 159±6 |
| | 4× | 145±0 |
| | 7× | 167±6 |
| | 10× | 167±4 |
| 50% MeOH | 2× | 173±2 |
| | 3× | 173±3 |
| | 4× | 176±6 |
| | 7× | 175±3 |
| | 10× | 188±6 |
| 100% ACN | 2× | 167±2 |
| | 3× | 167±5 |
| | 4× | 178±7 |
| | 7× | 184±13 |
| | 10× | 172±8 |
| 66.7% ACN | 2× | 183±4 |
| | 3× | 179±5 |
| | 4× | 186±19 |
| | 7× | 161±3 |
| | 10× | 171±2 |
| 50% ACN | 2× | 172±8 |
| | 3× | 185±6 |
| | 4× | 171±4 |
| | 7× | 194±7 |
| | 10× | 192±2 |

**Table S2. The number of identified metabolites from serum by different extraction solvent composition and volume.**

| Protein precipitation solvent composition | Extraction solvent volume | Identification number |
|---|---|---|
| 100% MeOH | 2× | 213±6 |
| | 3× | 230±2 |
| | 4× | 225±7 |
| | 7× | 229±2 |
| | 10× | 234±5 |
| 66.7% MeOH | 2× | 223±3 |
| | 3× | 213±1 |
| | 4× | 226±3 |
| | 7× | 212±3 |
| | 10× | 208±9 |
| 50% MeOH | 2× | 209±2 |
| | 3× | 198±1 |
| | 4× | 189±4 |
| | 7× | 192±2 |
| | 10× | 202±6 |
| 100% ACN | 2× | 169±7 |
| | 3× | 187±1 |
| | 4× | 186±6 |
| | 7× | 166±2 |
| | 10× | 174±2 |
| 66.7% ACN | 2× | 196±3 |
| | 3× | 173±1 |
| | 4× | 188±7 |
| | 7× | 172±4 |
| | 10× | 175±4 |
| 50% ACN | 2× | 211±8 |
| | 3× | 191±1 |
| | 4× | 220±13 |
| | 7× | 202±2 |
| | 10× | 214±3 |

**Table S3. The detailed information of the blastocysts corresponding to 120 SECM samples.**

| Biopsy results | Group | Biopsy days | Morphological grading |
|---|---|---|---|
| 45,XX,-4 | Aneuploid | D6 | 5BB (Fair) |
| 45,XN,-17 | Aneuploid | D6 | 6BC (Poor) |
| 45,XY,-5 | Aneuploid | D6 | 6BB (Fair) |
| 44,XN,-4,-18 | Aneuploid | D6 | 5BC (Poor) |
| 45,XN,-19 | Aneuploid | D7 | 6CB (Poor) |
| 45,XY,-21 | Aneuploid | D6 | 6AA (Good) |
| 45,XY,-13 | Aneuploid | D6 | 6BC (Poor) |
| 45,XN,-15 | Aneuploid | D5.5 | 5BB (Fair) |
| 45,XY,-15 | Aneuploid | D6.5 | 6BC (Poor) |
| 44,XX,-8,-13 | Aneuploid | D6 | 6BC (Poor) |
| 45,XN,-7 | Aneuploid | D6 | 4BB (Fair) |
| 45,XY,-22 | Aneuploid | D6.5 | 5BB (Fair) |
| 45,XY,-18 | Aneuploid | D6 | 6BB (Fair) |
| 45,XN,-11 | Aneuploid | D5.5 | 5BB (Fair) |
| 45,XN,-18 | Aneuploid | D6.5 | 5BB (Fair) |
| 45,XN,-19 | Aneuploid | D7 | 4BC (Poor) |
| 44,XN,-5,-16 | Aneuploid | D6 | 4BC (Poor) |
| 47,XX,+16 | Aneuploid | D5.5 | 4BB (Fair) |
| 47,XX,+16 | Aneuploid | D6 | 5BB (Fair) |
| 47,XX,+15 | Aneuploid | D5.5 | 5BB (Fair) |
| 47,XN,+14 | Aneuploid | D6.5 | 4AC (Poor) |
| 47,XN,+8 | Aneuploid | D6 | 5BC (Poor) |
| 45,XX,-10 | Aneuploid | D6 | 5BB (Fair) |
| 47,XY,+22 | Aneuploid | D6 | 5BB (Fair) |
| 48,XN,+18,+19 | Aneuploid | D6 | 6BB (Fair) |
| 47,XN,+11 | Aneuploid | D6 | 4BB (Fair) |
| 48,XY,+15,+22 | Aneuploid | D6 | 5BB (Fair) |
| 47,XN,+12 | Aneuploid | D7 | 6BC (Poor) |
| 47,XN,+13 | Aneuploid | D5.5 | 5BB (Fair) |
| 47,XY,+22 | Aneuploid | D6 | 6BB (Fair) |
| 47,XN,+21 | Aneuploid | D6 | 6AB (Good) |
| 47,XY,+19 | Aneuploid | D6 | 6AB (Good) |
| 47,XY,+19 | Aneuploid | D6 | 6AB (Good) |
| 45,XN,-16 | Aneuploid | D6 | 6BB (Fair) |
| 47,XX,+15 | Aneuploid | D6 | 5AA (Good) |
| 47,XN,+21 | Aneuploid | D5.5 | 5BB (Fair) |
| 48,XN,+11,+18 | Aneuploid | D6 | 6AB (Good) |
| 47,XX,+19 | Aneuploid | D6 | 6BC (Poor) |
| 48,XX,+3,+4 | Aneuploid | D6 | 5BB (Fair) |
| 46,XN,+19 | Aneuploid | D7 | 6BC (Poor) |
| 46,XO,+15 | Aneuploid | D6 | 4BC (Poor) |
| 47,XN,+11 | Aneuploid | D6 | 5AB (Good) |
| 48,XN,+20,+22 | Aneuploid | D6 | 5BB (Fair) |

| | | | |
|---|---|---|---|
| 45,XY,-14 | Aneuploid | D6 | 5BC (Poor) |
| 45,XN,-16 | Aneuploid | D6 | 5BC (Poor) |
| 45,XY,-6 | Aneuploid | D6 | 4BB (Fair) |
| 44,XX,-13,-15 | Aneuploid | D6 | 6BC (Poor) |
| 44,XN,-7,-21 | Aneuploid | D6 | 5BC (Poor) |
| 46,XN | Euploid | D6 | 5BB (Fair) |
| 46,XN | Euploid | D6 | 6BB (Fair) |
| 46,XN | Euploid | D6 | 6BB (Fair) |
| 46,XN | Euploid | D6 | 6BB (Fair) |
| 46,XN | Euploid | D6 | 6BB (Fair) |
| 46,XN | Euploid | D6 | 6BB (Fair) |
| 46,XN | Euploid | D6 | 6BB (Fair) |
| 46,XN | Euploid | D6 | 6BB (Fair) |
| 46,XN | Euploid | D6 | 4BB (Fair) |
| 46,XN | Euploid | D6 | 4BB (Fair) |
| 46,XN | Euploid | D5.5 | 4BB (Fair) |
| 46,XN | Euploid | D6 | 5BB (Fair) |
| 46,XN | Euploid | D5.5 | 4BB (Fair) |
| 46,XN | Euploid | D6 | 4BB (Fair) |
| 46,XN | Euploid | D6 | 4BB (Fair) |
| 46,XN | Euploid | D5.5 | 5BB (Fair) |
| 46,XN | Euploid | D5.5 | 5BB (Fair) |
| 46,XN | Euploid | D6 | 5BB (Fair) |
| 46,XN | Euploid | D6 | 6BB (Fair) |
| 46,XN | Euploid | D5.5 | 5BB (Fair) |
| 46,XN | Euploid | D5.5 | 5BB (Fair) |
| 46,XN | Euploid | D5.5 | 5BB (Fair) |
| 46,XN | Euploid | D6 | 6BB (Fair) |
| 46,XN | Euploid | D6 | 5BB (Fair) |
| 46,XN | Euploid | D6 | 5BA (Good) |
| 46,XN | Euploid | D6 | 5AB (Good) |
| 46,XN | Euploid | D6 | 6AA (Good) |
| 46,XN | Euploid | D6 | 6AA (Good) |
| 46,XN | Euploid | D6 | 6AA (Good) |
| 46,XN | Euploid | D6 | 6AA (Good) |
| 46,XN | Euploid | D6 | 6AB (Good) |
| 46,XN | Euploid | D6 | 6AB (Good) |
| 46,XN | Euploid | D6 | 6AB (Good) |
| 46,XN | Euploid | D6 | 5BA (Good) |
| 46,XN | Euploid | D6 | 6AB (Good) |
| 46,XN | Euploid | D5.5 | 5BA (Good) |
| 46,XN | Euploid | D6 | 5BA (Good) |
| 46,XN | Euploid | D6 | 5AB (Good) |
| 46,XN | Euploid | D6 | 5AB (Good) |
| 46,XN | Euploid | D5.5 | 5AB (Good) |
| 46,XN | Euploid | D5.5 | 6AA (Good) |

| | | | |
|---|---|---|---|
| 46,XN | Euploid | D6 | 6AB (Good) |
| 46,XN | Euploid | D6 | 5BA (Good) |
| 46,XN | Euploid | D6 | 6BA (Good) |
| 46,XN | Euploid | D6 | 6BA (Good) |
| 46,XN | Euploid | D6 | 6BA (Good) |
| 46,XN | Euploid | D6 | 6BA (Good) |
| 46,XN | Euploid | D5.5 | 5AB (Good) |
| 46,XN | Euploid | D6 | 6BC (Poor) |
| 46,XN | Euploid | D6 | 6BC (Poor) |
| 46,XN | Euploid | D6 | 6BC (Poor) |
| 46,XN | Euploid | D6 | 5BC (Poor) |
| 46,XN | Euploid | D6.5 | 5BC (Poor) |
| 46,XN | Euploid | D6 | 5BC (Poor) |
| 46,XN | Euploid | D6 | 5BC (Poor) |
| 46,XN | Euploid | D5.5 | 5BC (Poor) |
| 46,XN | Euploid | D6 | 5BC (Poor) |
| 46,XN | Euploid | D6 | 6BC (Poor) |
| 46,XN | Euploid | D6 | 6BC (Poor) |
| 46,XN | Euploid | D7 | 6BC (Poor) |
| 46,XN | Euploid | D6 | 6BC (Poor) |
| 46,XN | Euploid | D6 | 4BC (Poor) |
| 46,XN | Euploid | D6 | 5BC (Poor) |
| 46,XN | Euploid | D6 | 5BC (Poor) |
| 46,XN | Euploid | D6 | 6AC (Poor) |
| 46,XN | Euploid | D6 | 6BC (Poor) |
| 46,XN | Euploid | D7 | 6BC (Poor) |
| 46,XN | Euploid | D6 | 6BC (Poor) |
| 46,XN | Euploid | D6 | 6BC (Poor) |
| 46,XN | Euploid | D6 | 6BC (Poor) |
| 46,XN | Euploid | D6 | 6BC (Poor) |
| 46,XN | Euploid | D6 | 6BC (Poor) |

**Table S4. The fifteen candidate biomarkers preselected based on univariate and multivariate statistical criteria and analytical stability (|FC| > 1.2, p < 0.05, CV< 0.25).**

| Compound | Fold Change(E vs. A) | *p* value | CV |
|---|---|---|---|
| 1,5-Naphthalenediamine | 0.8239 | 3.50E-08 | 0.1528 |
| 12-Aminododecanoic acid | 0.3942 | 3.46E-03 | 0.2044 |
| 1-Phenyl-1H-pyrazolo[3,4-d]pyrimidin-4-amine | 1.4212 | 1.87E-06 | 0.2174 |
| 2,4-Dinitrophenol | 1.2170 | 8.09E-06 | 0.1423 |
| Adrenic acid | 1.2615 | 4.63E-05 | 0.2441 |
| Corticosterone | 1.4027 | 3.34E-02 | 0.2036 |
| DL-Tryptophan | 1.2943 | 1.10E-09 | 0.2238 |
| Docosapentaenoic acid | 1.2342 | 6.74E-06 | 0.2112 |
| Dodecanedioic acid | 1.2075 | 4.70E-06 | 0.2079 |
| Dodecylamine | 1.4325 | 2.26E-08 | 0.2062 |
| Dodecyltrimethylammonium | 1.2953 | 2.63E-08 | 0.2048 |
| Hexadecanamide | 2.0877 | 8.03E-08 | 0.1987 |
| Mono(2-ethylhexyl) phthalate (MEHP) | 1.2123 | 7.81E-03 | 0.2454 |
| Oleoyl ethanolamide | 1.2004 | 9.57E-03 | 0.1499 |
| trans-3-Indoleacrylic acid | 1.2528 | 1.00E-07 | 0.2165 |

**Table S5. Performance metrics for each model, including accuracy, sensitivity, specificity, PPV, NPV, AUC, and F1 score.**

| Model | Sensitivity | Specificity | PPV | NPV | Accuracy | F1 Score | AUC |
|---|---|---|---|---|---|---|---|
| Logistic Regression | 0.9167 | 0.9583 | 0.9362 | 0.9452 | 0.9417 | 0.9263 | 0.9659 |
| LDA | 0.8750 | 0.9583 | 0.9333 | 0.9200 | 0.9250 | 0.9032 | 0.9737 |
| SVM | 0.8958 | 0.9583 | 0.9348 | 0.9324 | 0.9333 | 0.9149 | 0.9630 |
| Random Forest | 0.9583 | 0.9861 | 0.9787 | 0.9726 | 0.9750 | 0.9684 | 0.9861 |
| KNN | 0.9375 | 0.8750 | 0.8333 | 0.9545 | 0.9000 | 0.8824 | 0.9576 |
| Decision Tree | 0.8750 | 0.8611 | 0.8077 | 0.9118 | 0.8667 | 0.8400 | 0.7975 |
| Naïve Bayes | 0.8750 | 0.8194 | 0.7636 | 0.9077 | 0.8417 | 0.8155 | 0.9392 |
| Ensemble | 0.8958 | 1.0000 | 1.0000 | 0.9351 | 0.9583 | 0.9451 | 0.9766 |

**Table S6. Twelve candidate biomarkers preselected based on univariate and multivariate statistical criteria and analytical stability (|FC| > 1.2, p < 0.05, CV< 0.15).**

| Compound | G vs. F | | G vs. P | | F vs. P | | CV |
|---|---|---|---|---|---|---|---|
| | Fold Change (G vs. F) | *p* value | Fold Change (G vs. P) | *p* value | Fold Change (F vs. P) | *p* value | |
| 1-Tetradecylamine | 0.9069 | 1.32E-01 | 0.6079 | 1.46E-04 | 0.6703 | 4.50E-04 | 0.1500 |
| 2,2,6,6-Tetramethyl-1-piperidinol (TEMPO) | 0.8553 | 5.40E-02 | 0.8254 | 2.40E-02 | 0.9649 | 6.58E-01 | 0.1153 |
| 2-hexyl-5-(4-nonylphenyl)pyrimidine | 0.9101 | 4.04E-01 | 0.6687 | 6.01E-07 | 0.7347 | 1.48E-06 | 0.1467 |
| 2-Hydroxyvaleric acid | 1.0813 | 2.10E-01 | 1.2010 | 6.25E-05 | 1.1107 | 3.09E-02 | 0.0861 |
| 4-Hydroxycoumarin | 0.7403 | 2.43E-03 | 0.7649 | 3.58E-04 | 1.0331 | 6.81E-01 | 0.1265 |
| Decanoylcarnitine | 0.7821 | 7.75E-05 | 0.7851 | 1.21E-03 | 1.0038 | 9.51E-01 | 0.1393 |
| Milrinone | 1.6194 | 4.92E-02 | 1.0332 | 8.84E-01 | 0.6380 | 2.11E-02 | 0.1032 |
| N,N-Diisopropylethylamine (DIPEA) | 0.7894 | 3.73E-02 | 0.8038 | 3.60E-02 | 1.0183 | 8.52E-01 | 0.1221 |
| N,N-Dimethyldecylamine N-oxide | 1.1460 | 1.72E-01 | 0.5525 | 1.60E-01 | 0.4821 | 4.19E-02 | 0.1473 |
| Oleoyl ethanolamide | 0.8232 | 1.69E-01 | 0.8031 | 3.17E-02 | 0.9756 | 7.73E-01 | 0.1499 |
| Proline | 1.1357 | 3.89E-01 | 1.3066 | 3.34E-02 | 1.1505 | 1.87E-01 | 0.0449 |
| Rabenzazole | 1.1293 | 1.31E-02 | 1.2365 | 3.87E-04 | 1.0949 | 8.45E-02 | 0.1308 |

**Table S7. Performance metrics for each model, including overall accuracy, Kappa coefficient, macro-average sensitivity, specificity, PPV, NPV, F1 score, and AUC.**

| Model | Overall Accuracy | Kappa | Sensitivity | Specificity | PPV | NPV | F1 Score | AUC |
|---|---|---|---|---|---|---|---|---|
| Multinomial Regression | 0.7361 | 0.6042 | 0.7361 | 0.8681 | 0.7348 | 0.8706 | 0.733 | 0.8825 |
| LDA | 0.8056 | 0.7083 | 0.8056 | 0.9028 | 0.8149 | 0.9049 | 0.8049 | 0.9039 |
| SVM | 0.7639 | 0.6458 | 0.7639 | 0.8819 | 0.7804 | 0.8855 | 0.7621 | 0.91 |
| Random Forest | 0.7778 | 0.6667 | 0.7778 | 0.8889 | 0.7787 | 0.8903 | 0.7763 | 0.9184 |
| KNN | 0.7083 | 0.5625 | 0.7083 | 0.8542 | 0.7039 | 0.8566 | 0.7047 | 0.8786 |
| Decision Tree | 0.625 | 0.4375 | 0.625 | 0.8125 | 0.6221 | 0.8141 | 0.6221 | 0.7075 |
| Naive Bayes | 0.6111 | 0.4167 | 0.6111 | 0.8056 | 0.6278 | 0.8117 | 0.6023 | 0.8368 |
| Ensemble | 0.8056 | 0.7083 | 0.8056 | 0.9028 | 0.8083 | 0.9036 | 0.8052 | 0.9253 |

**Table S8. Class-specific performance metrics, including sensitivity, specificity, PPV, NPV, F1 score and AUC.**

| Group | Sensitivity | Specificity | PPV | NPV | F1 Score | AUC |
|---|---|---|---|---|---|---|
| G | 0.8750 | 0.8958 | 0.8077 | 0.9348 | 0.8400 | 0.9444 |
| F | 0.7917 | 0.8750 | 0.7600 | 0.8936 | 0.7755 | 0.8889 |
| P | 0.7500 | 0.9375 | 0.8571 | 0.8824 | 0.8000 | 0.9427 |

#### References


1. Wishart, D. S.; Guo, A.; Oler, E.; Wang, F.; Anjum, A.; Peters, H.; Dizon, R.; Sayeeda, Z.; Tian, S.; Lee, B. L.; Berjanskii, M.; Mah, R.; Yamamoto, M.; Jovel, J.; Torres-Calzada, C.; Hiebert-Giesbrecht, M.; Lui, V. W.; Varshavi, D.; Varshavi, D.; Allen, D.; Arndt, D.; Khetarpal, N.; Sivakumaran, A.; Harford, K.; Sanford, S.; Yee, K.; Cao, X.; Budinski, Z.; Liigand, J.; Zhang, L.; Zheng, J.; Mandal, R.; Karu, N.; Dambrova, M.; Schioth, H. B.; Greiner, R.; Gautam, V., HMDB 5.0: the Human Metabolome Database for 2022. *Nucleic Acids Res* **2022,** *50* (D1), D622-D631.

2. Kim, S.; Chen, J.; Cheng, T.; Gindulyte, A.; He, J.; He, S.; Li, Q.; Shoemaker, B. A.; Thiessen, P. A.; Yu, B.; Zaslavsky, L.; Zhang, J.; Bolton, E. E., PubChem 2025 update. *Nucleic Acids Res* **2025,** *53* (D1), D1516-D1525.

3. Pang, Z.; Chong, J.; Zhou, G.; de Lima Morais, D. A.; Chang, L.; Barrette, M.; Gauthier, C.; Jacques, P. E.; Li, S.; Xia, J., MetaboAnalyst 5.0: narrowing the gap between raw spectra and functional insights. *Nucleic Acids Res* **2021,** *49* (W1), W388-W396.
